# Parental fasting effects on offspring immune gene expression and gut microbiota in a species with male pregnancy (*Syngnathus typhle)*

**DOI:** 10.1101/2025.06.30.662305

**Authors:** Freya A. Pappert, Nils Newrzella, Olivia Roth

## Abstract

Intermittent fasting is widely promoted for its potential to improve health and extend lifespan, yet these benefits may come at a reproductive cost, potentially reducing parental fitness and offspring quality. While the inter- and transgenerational effects of fasting are increasingly studied, they remain poorly understood in species with unconventional reproductive roles. Investigating such effects in these systems is crucial, as the evolutionary trade-offs between somatic maintenance and reproductive investment may differ from those in species with conventional reproductive roles. In this study, we investigated the effects of intermittent fasting (IF) in a species with male pregnancy, *Syngnathus typhle*, by exposing mothers and fathers to either IF or ad libitum (AL) feeding before mating. Upon transfer of maternal eggs to paternal brood pouches, males remained on their assigned diets throughout pregnancy. Offspring from all parental diet combinations AL(p)xAL(m), IF(p)xIF(m), AL(p)xIF(m), IF(p)xAL(m) were analyzed at birth before first feeding alongside parents for morphology, immune and epigenetic candidate gene expression, and gut microbiota composition. Mothers under IF showed greater condition loss, leading to reduced offspring condition regardless of paternal diet, highlighting the importance of maternal provisioning through eggs. However, IF fathers exhibited increased immune activation and microbiome shifts that were mirrored in offspring, suggesting paternal priming via epigenetic and microbial inheritance. Offspring from mismatched parental diets showed disrupted immune-microbiome correlations, indicating that aligned parental cues support more stable offspring development. These findings highlight how parental nutritional history shapes offspring resilience and emphasize the value of studying species with diverse reproductive strategies and life histories to understand the full spectrum of trans-generational plasticity in nature.

## 1. Introduction

An organism’s traits are shaped not only by the genes it inherits, but also by non-genetic influences from the environment (within-generational phenotypic plasticity, WGP), as well as the conditions the parents experienced (trans-generational plasticity, TGP) (1–3). DNA-independent inheritance can occur through multiple, often overlapping, routes including reproductive elements such as eggs (and to a lesser extent sperm), as well as through parental care, feeding, or other forms of close contact. These routes transmit a diverse array of information: for instance, eggs may carry maternal hormones and metabolites (4–6), epigenetic marks shaped by parental life history (7,8), immunological components such as antibodies (i.e., transgenerational immune priming - TGIP) (9–12), and even vertically inherited microbial communities (13–15). In contrast, postnatal influences such as parental care or feeding may shape offspring phenotype through continued microbial transfer, behavioral imprinting, or immune priming (16). Understanding the relative contributions of these factors to the expression of offspring phenotypes and ultimately their fitness, and how they change over time, is crucial for predicting how organisms acclimatize, and ultimately adapt, to new and changing environments (1). While plasticity, including TGP, can buffer organisms against stress in the short term, it can also generate phenotypic variation, some of which may confer a fitness advantage under novel conditions and thus be favored by selection, potentially buying time for genetic changes to follow (17,18).

When mothers experience changes in their environment, this can alter offspring development via maternal effects, even in the absence of direct exposure during the offspring’s own early experiences (1). For example, a child may grow up in a food-rich environment but exhibit traits shaped by maternal exposure to food scarcity (e.g., increased fat deposition). These traits could become beneficial if the child later encounters similar conditions. Since the mother has already navigated a famine or nutrient scarcity, she may pass down helpful cues to prepare her offspring for similar challenges (19). In some cases, phenotypic adjustment in response to current environmental cues (WGP) may offer a better match to local conditions than relying on parental cues. However, when rapid or early-life responses are required, offspring may benefit from parental pre-conditioning via TGP (1). A well-documented example of the phenomenon of gestational programming is the development of a “thrifty phenotype” in offspring of mothers who faced nutritional constraints during pregnancy. Offspring of mothers with intrauterine growth restriction (IUGR) often have low birth weight and are predisposed to metabolic diseases like obesity, hypertension, and diabetes later in life (20–23). These offspring are programmed to efficiently acquire and use nutrients, a trait advantageous in environments with scarce food. However, in nutrient-rich settings, such acclimatization can lead to negative maladaptive outcomes like obesity (19). Across some mammalian species, including rats and sheep (24–26), studies have shown similar links between maternal nutrient deficiencies during pregnancy and adult health issues in the offspring, further highlighting the broad impact of maternal experiences on offspring development (19).

Paternal effects are often overlooked in transgenerational research, partly because fathers do not gestate offspring and may have limited parental involvement. However, growing evidence suggests that fathers play a significant role in inheritance beyond genetics, particularly through sperm-mediated effects such as epigenetic modifications and small RNAs (27). Advances in epigenetics are shifting the research landscape to recognize these paternal contributions. In parallel, genetic factors such as increased germline mutations with paternal age can also affect offspring health and lifespan by reducing sperm quality and integrity (28,29). Age-related changes can also affect resource allocation to ejaculate (30,31), and in some species, offspring of older fathers have shorter lifespans and a higher incidence of chronic illnesses (32,33), likely resulting from a combination of genetic and phenotypic changes associated with aging and cumulative environmental stress (34). Beyond age, paternal diet can also shape offspring health, influencing metabolic function and early developmental outcomes (35). Studies have shown that exposure to a high-fat diet before mating can alter offspring traits, including body weight and metabolic parameters, though these effects vary across species (36,37). These findings underscore the importance of considering paternal trans-generational plastic effects in shaping offspring development (35).

Although parental effects have been studied across diverse taxa, including insect and fish (9,38,39), many models of trans-generational plasticity focus on species with conventional reproductive roles, where females typically both gestate and provide the majority of parental care. As a result, we still lack a clear understanding of how both parents’ environmental experiences - before and during reproduction - contribute to offspring phenotype, particularly in species with unconventional reproductive strategies. Teleost fish exhibit a remarkable range of variation in parental care strategies. Among those species that do exhibit parental care, approximately half provide exclusive paternal care (40,41). In particular, the Syngnathidae family is renowned for its exceptionally dedicated male parental care, having evolved a range of male brooding forms (42,43). These adaptations span from the simple attachment of eggs to the male’s ventral surface (e.g., *Nerophis ophidion*) to the development of highly specialized brooding structures (e.g., *Hippocampus erectus*). These brooding structures are analogous to those found in eutherian mammals, and they can provide the developing embryos with protection, nutrients, oxygen, and immunological components (9,44–47). Furthermore, they can mediate transgenerational effects in response to environmental conditions such as salinity levels affecting offspring gene expression and survival (48), and shape the external juvenile microbiome through paternal provisioning (49). Remarkably, the phenomenon of male pregnancy has its origins in a single evolutionary event, which then gave rise to approximately 300 distinct species of syngnathid (42). The Syngnathidae family can be found in a broad range of temperate and tropical environments, including rocky and coral reefs, mangrove forests, seagrass beds, estuaries, and rivers (42).

Species like the broadnosed pipefish (*Syngnathus typhle*), with their reversed reproductive roles and male pregnancy, offer a unique opportunity to disentangle the contributions of maternal egg production from the physiological changes associated with paternal pregnancy (e.g., body reshaping and energy allocation) (45). In this pipefish species, females typically produce on average more eggs than males can brood (50,51), making males the more reproductively constrained sex (52). Due to their polygamous mating system and the males’ limited reproductive capacity, males become the more choosy sex, often preferring larger females, while females exhibit ornamentation and actively court males (50,52–55). This species’ non-conventional mating system and male pregnancy make it an ideal model for studying how parental effects and resource allocation are shaped by sex-specific reproductive strategies. It allows for a more nuanced understanding of how males and females may influence offspring development differently, especially when disentangling gestational vs. sex-specific effects on offspring phenotype (56–58).

In this study we examined the influence of dietary restriction in *Syngnathus typhle* by subjecting both males and females to intermittent fasting (IF) for a month prior to reproduction. Specifically, the experimental conditions included a twice-daily feeding schedule with live food for ad libitum (AL) conditions, or IF with a feeding day followed by two fasting days. Individuals were then mated following various dietary combinations, with paternal (p) vs. maternal (m) combinations, hereafter abbreviated as AL (p) x AL (m), IF (p) x IF (m), AL (p) x IF (m), and IF (p) x AL (m). The IF treatment was continued for males throughout their pregnancy. After offspring birth, the young were sacrificed before first feeding alongside their parents to assess morphological characteristics, immune- and epigenetic-related gene expression, and gut microbiota composition through 16S rRNA sequencing. We hypothesized that offspring from IF parents would be morphologically smaller than those from AL parents, due to limited parental resource availability. Because males gestate the offspring in *Syngnathus typhle*, we expect paternal diet to have a stronger influence on offspring traits. In particular, that the offspring’s gut microbiota and immune-related gene expression would reflect paternal feeding conditions, consistent with transgenerational plasticity mediated by male pregnancy.

Our findings open new avenues to explore how parental diet and sex-specific reproductive roles influence offspring development, particularly in species with reversed reproductive roles and male pregnancy. This study provides critical insights into transgenerational effects, gestational influences, and the role of the microbiome in offspring health, while challenging existing paradigms by considering both maternal and paternal effects in shaping offspring phenotype.

## 2. Material and methods

### 2.1 Study species and collection area

*Syngnathus typhle* were caught in the southwest Baltic Sea on the island Fehrman at the bay of Orth (54° 272 N, 11° 32 O) late April of 2022, using snorkelling gear and hand nets. The broadnosed pipefish were then transported back to Kiel University in appropriate containers connected to air stones filled with Baltic Sea Water from the field. At the institute, they were divided into sex-aggregated quarantine tanks maintained at 18 ppt salinity and 12°C. To prevent the introduction of harmful parasites into the aquaria, the fish received three, one-hour anti-parasite treatments using a 37% formalin solution at a 1:8000 concentration. Subsequently, the fish were allocated to sex-disaggregated 100-liter tanks (50Lx40Wx51D), where they were gradually acclimated over two weeks to their new environment. During this period, they were transitioned from live food to frozen mysids. The final water temperature was maintained at 18 °C, and fish were kept under a simulated day-night light cycle (i.e., 14L:10D).

### 2.2 Experimental design

Pipefish were tagged with elastomer implants in red, blue, or orange, by lightly sedating them with 0.02% Tricaine mesylate (MS-222, Sigma-Aldrich). Tags were injected subcutaneously using a syringe on the left side of the body (Figure S1). During this brief procedure, the fish were also measured for size and weight.

The tagged fish were then sorted into groups of three (one red, one blue, and one orange), separated by sex, and housed in 28 aquaria. Each aquarium was randomly assigned to either ad libitum (AL) food or intermittent fasting (IF) treatment (total n = 84: 21 AL males, 21 AL females, 21 IF males, 21 IF females). Fish in the AL treatment were fed defrosted mysids twice daily (8 a.m. and 4 p.m.), while those in the IF treatment were fed every third day, leading to two consecutive days of fasting. Given that *Syngnathus typhle* lack stomachs (59), overfeeding was unlikely; thus, the fasting group experienced a caloric reduction. To ensure observed effects were due to fasting and not nutrient deficiencies, both groups received enriched mysids every 6th day. The food was supplemented with multivitamins (15 µL of JBL Atvitol) and fatty acids (Omega-3 1000 capsules), allowed to absorb into the thawed mysids for 2-3 minutes. The feeding regime lasted 30 days, during which the water temperature was gradually increased by 1°C every five days to reach 18°C, simulating the natural conditions of the Baltic Sea known to trigger optimal mating behaviour in pipefish.

After the month-long treatment, the pipefish were removed from their respective aquaria and paired for mating, creating the following crosses (each replicated 7 times): AL (p) × AL (m), IF (p) × IF (m), AL (p) × IF (m), and IF (p) × AL (m). A mating window of 24-48 hours was provided, which is typically sufficient for successful mating, as both sexes had been separated for a month and were primed to mate at 18°C. Pregnant males were returned to their treatment tanks, while females were euthanized with an overdose of MS-222 (500 mg/L), measured for total body length and weight, and dissected for further analysis (Table S1). The pregnant males were kept on their AL or IF diet and approximately one month later, they gave birth. At the time of birth, five offspring (or as many as were present) were collected from each male. Both the fathers and offspring were euthanized, measured, and dissected. The initial tagging and tank assignments allowed for accurate matching of each fish to its initial size and weight measurements.

### 2.3 Sample collection

After euthanasia, parental pipefish were dissected to remove the head kidney and liver, which were placed into 2 mL Eppendorf tubes containing RNAlater. These samples were stored in a refrigerator for two days and then transferred to -20°C awaiting further analysis. The intestinal tract was also removed, and a 1 cm section from the aboral part was placed into a sterile 1.5 mL Eppendorf tube using sterile tools. This sample was flash-frozen and stored at -80°C. All remaining gut tissue and gonads were collected for fat analysis.

Due to their small size, offspring were measured using a transparent container positioned over millimeter paper, and images were analyzed for size using ImageJ (Version 1.53t; Schneider, Rasband, and Eliceiri 2012) (Figure S2). Offspring were then euthanized and weighed using a precision scale. The gut was delicately extracted and placed into individual PCR tubes, which were stored at -80°C. The remaining body was submerged in RNAlater and processed like the adult tissue samples.

### 2.4 Fat content measurement

Parent fat content was measured following the methods described by Auer (2010) (60). The tissues were dried in an oven for 2 days at 60°C and weighed using a precision scale. Samples were then submerged in anhydrous diethyl ether to dissolve triglycerides for 30 minutes, after which the ether (containing the dissolved fat) was discarded. The samples were air-dried for 1 hour and weighed again. This process was repeated until the samples reached a constant lean dry weight. The total fat content was then measured by subtracting the initial weight with the final dried weight (Table S1).

### 2.5 RNA extraction, reverse transcription and gene expression

We quantified the mRNA levels of 44 pre-selected target genes based on previous studies (9,61,62) using quantitative real-time polymerase chain reaction (qRT-PCR). This was done on a 96.96 dynamic array BioMark HD system (GE chip; Fluidigm, South San Francisco, CA, USA) as described by Beemelmanns & Roth (2016) (9). The selected genes are listed in Table 1 and supplementary material Table S3. These genes were grouped into functional categories, following previous literature: (i) innate immune system (immediate, non-specific immune defense, e.g., phagocytosis), (ii) adaptive immune system (specific, antibody-mediated immune defense), (iii) innate and adaptive immune genes (genes involved in both pathways), (iv) complement system (supporting antibody and phagocytic cell-mediated responses), and (v) epigenetic modulators (genes involved in DNA methylation, histone de/methylation, and histone de/acetylation) (Tables S3).

**Table 1:**
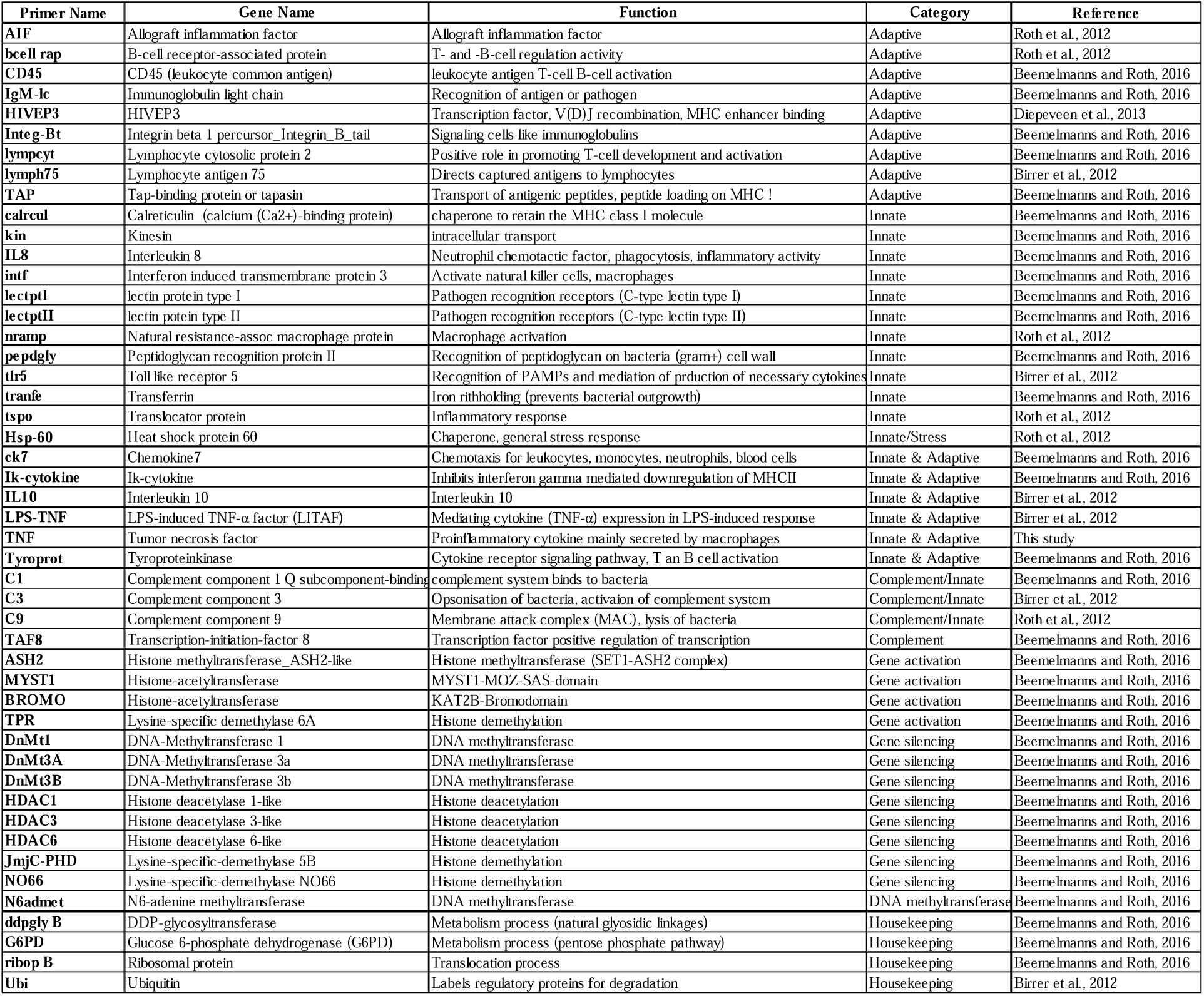
Name and function of 44 target genes and 4 housekeeping genes. Listed are all genes from different functional categories: adaptive, innate, innate & adaptive, complement system, gene activation, gene silencing

RNA was extracted from the head kidneys of successfully mated males and females, as well as from whole-body tissue samples of five offspring from each parent, using the RNeasy Mini Kit (Qiagen, Venlo, Netherlands). The RNA concentration was measured with a DeNovix DS-11 Nanodrop, and all samples were normalized to 50 ng/µL before being stored at -80°C. cDNA synthesis was performed using the RevertAid First Strand cDNA Synthesis Kit (Thermo Scientific, K1621) according to the manufacturer’s protocol, and stored again at -80°C for later use.

A pre-amplification step was conducted by mixing 1 uL of each of the 48 primers (500 nM, forward and reverse) into a primer pool. The reaction mixture consisted of 264 uL 2X TaqMan PreAmp Master Mix (Applied Biosystems, Waltham, MA, USA), 52.8 uL primer pool mix, 79.2 uL purified water. Then 3.8 uL fo the mixture was combined with 1.3 uL of the cDNA sample. A dilution series (1:10, 1:20, 1:40, 1:80, 1:160, 1:320) was prepared to validate primer efficiencies (Table S3). The PCR settings were: 95°C for 10 min, followed by 14 cycles of 95°C for 15 sec and 60°C for 4 min. The resulting PCR products were diluted 1:10 with low EDTA-TE buffer (10 mM Tris, 0.1 mM EDTA, pH 8) (63).

For the sample pre-mix, we combined 369.6 uL Ssofast-EvaGreen Supermix with Low ROX (Bio-Rad Laboratories, Hercules, CA, USA) with 37 uL 20x DNA Binding Dye Sample & Assay Loading Reagent (Fluidigm). Each well of the 96-well plate was loaded with 3.9 uL of this sample pre-mix, followed by 3.1 uL of the diluted pre-amplified PCR products (in duplicates). The assay mix for the chip consisted of 369.6 uL Assay Loading Reagent (Fluidigm) and 295.7 uL low EDTA-TE buffer. A total of 6.3 uL of the assay mix was loaded into each well of a 96-well plate, and 0.7 uL of a 50 uM primer pair mix was added to each well (in duplicates). Finally, the GE chips were loaded with 5 uL of sample mix and 5 µL of assay mix, and run on the BioMark system using the GE-fast 96.96 PCR+Melt v2.pcl protocol (Fluidigm). The plates included no-template controls (NTC), controls for gDNA contamination (NEG), and a serial dilution as mentioned previously.

### 2.6 DNA extraction and sequencing

To extract microbiota from the adult hind-gut and the entire offspring gut, DNA was isolated using the DNeasy Blood & Tissue Kit (QIAGEN, Germany), following the manufacturer’s protocol with modifications for Gram-positive bacteria based on (64). A 16S PCR was then performed to confirm successful extraction. Library preparation for a total of 114 samples (including: 20 AL(p)xAL(m), 15 AL(p)xIF(m), 20 IF(p)xAL(m), 25 IF(p)xIF(m) offspring, along with 8 AL and 9 IF adult males, and 9 AL and 8 IF adult females) was completed by the Institute for Experimental Medicine (UKSH, Campus Kiel). Sequencing of the V3-V4 hypervariable region (341f/806r) was carried out using the Illumina MiSeq platform (Illumina, USA) with 2x 300-bp paired-end settings at IKMB Kiel.

### 2.7 Morphological and fat content data analysis

Statistical analyses were conducted using RStudio (Version 2024.04.2). We assessed changes in total body length (cm) and weight (g) before and after treatment for both male and female pipefish. Normality of the data was evaluated using the Shapiro-Wilk test, and given the small sample sizes, we also tested for homogeneity of variance with Levenes test (Table S2). To assess changes in body size over time (pre- and post-treatment), we used linear mixed-effects models, accounting for individual variation (e.g., lmer(Body_Length ∼ Time * treatment + sex + (1 | ID)); Table S2_morphology). This approach was also applied to the weight data. Since sex could introduce confounding effects, especially considering that females only underwent one month of treatment compared to two months for males, we also analyzed males and females separately. For these analyses, we performed one-way ANOVAs on initial and final length and weight, using treatment as a factor (e.g., aov(Start_length ∼ treatment; Table S2_morphology). Additionally, we calculated Fulton’s condition factor (weight/length³) to better assess overall fish condition (65), to which the same statistics were applied (Table S2_morphology). The fat body content was also measured for normality and variance distribution. After which a Kruskal-Wallis rank sum test with Benjamin-Hochberg correction was done for treatment and sex (Table S2_morphology). To test crossed parental dietary treatment effect for offspring size and weight we used a one-way anova (e.g.: weight ∼ crosstreatment) and if significant, we followed up with a TukeyHSD test (Table S2_morphology).

### 2.8 Differential gene expression analysis

The quality of the chip run output was assessed by reviewing the Chip Run Info (Table S4). Samples flagged as problematic, and negative controls, were removed prior to further analysis. Each sample was analysed in two technical replicates, and the mean cycle threshold (Ct) value was calculated for each sample. Primer efficiency values were determined from a dilution series for all primers (1:10, 1:20,1:40, 1:80, 1:160, 1:320). Any primer with efficiency below 80% was excluded from further analysis, including primers for the following genes: *C3*, *C9*, *Kin*, *ASH2*, *Ck7*, *N6admet*, *TAF8*, *Tyroprot*, and *tlr5* (Table S3). The geometric mean of Ct values for housekeeping genes (*ddpgly.B, G6PD, ribop.B*, and *Ubi*) was used to quantify relative gene expression for each target gene, calculated as ΔCt values (ΔCt = mean Ct of target gene - geometric mean Ct of housekeeping genes). Samples or genes with excessive missing values were also excluded (i.e., *TNF*, with 25 NA values, Table S4). This cleaning process resulted in a final dataset comprising 34 genes and 104 samples (i.e., two plate chips), including 19 offspring from AL(p)xAL(m), 22 offspring from IF(p)xIF(m), 16 offspring from AL(p)xIF(m), and 17 offspring from IF(p)xAL(m) treatments.

A principal component analysis (PCA) was conducted to investigate differential gene expression profiles across parental dietary treatments (using the prcomp function in R). Additionally, a PERMANOVA model (utilizing the adonis2 function in the vegan package; e.g., adonis2(pca$x[,1] ∼ Treatment * Family * Plate, permutations = 999, method = “bray”); Table S2_PermanovaPCA) was applied based on a Bray-Curtis distance matrix of the principal components. Separate PCA analyses were conducted for parent-only and offspring-only datasets to examine treatment effects independently for mothers, fathers, and offspring combinations (Table S2_PermanovaPCA). Ellipses represent 80% confidence intervals. To assess treatment effects on individual genes, linear mixed models (LMMs) were applied with plate included as a random effect (lmer(gene ∼ Treatment + (1|Plate), *lme4* package, Table S2_LMM). Family was included in the PCA PERMANOVA model but was not significant. Consequently, family was excluded from the final LMMs to avoid overfitting and model instability. This analysis was performed separately for males, females, and offspring. Genes that reached statistical significance were then mapped onto the PCA plot with their respective loadings. In this PCA visualization, arrow directions indicate gene correlations with the principal components, while arrow lengths represent each genès proportional contribution to overall variance.

To calculate relative fold changes (logFC), the ΔΔCt method was employed, representing the difference between the ΔCt values of treatment and control groups. For parental samples (mothers and fathers), ΔΔCt was calculated as ΔCt_IF - ΔCt_AL. For offspring, ΔΔCt values compared combinations of parental treatments (ΔCt_IF(p)xAL(m) - ΔCt_AL(p)xAL(m)). Relative gene expression was calculated as 2×(-ΔΔCt) and then log2-transformed to normalize the distribution.

Interaction plots were created to illustrate gene expression patterns as influenced by maternal or paternal diet. These plots display mean ΔCt values (multiplied by -1 for directionality) with error bars representing the standard error.

### 2.9 Microbiome data analysis

The 16S rRNA amplicon sequencing analysis was carried out on the QIIME2 platform (v.2022.8.3) (66). Using dada2, raw paired-end Illumina reads were initially demultiplexed, with primers for the 16S V3-V4 region then trimmed. Quality control involved denoising to distinguish genuine sequence diversity from sequencing errors. Further filtering removed low-quality reads (trimming forward reads to 260 bp and reverse reads to 210 bp). Forward and reverse reads were merged, and chimeric sequences were removed. A phylogenetic tree was constructed using FastTree (v.2.1.11) based on approximately-maximum-likelihood from the longest root (67). Interactive α rarefaction curves were generated at a max depth of 20,000, assessing if sequencing depth was sufficient to capture the full bacterial community. Taxonomic classification for the V3/V4 hypervariable region was done with the SILVA v138 database and a naive Bayes classifier (68). Amplicon sequence variants (ASVs) were filtered to exclude chloroplast and mitochondrial sequences and were exported at the genus level as operational taxonomic units (OTUs) for further analysis in RStudio (v.2022.07.2).

The data was then filtered by removing singletons and applying a threshold of 0.2% of total reads per sample. After filtering, a total of 189 operational taxonomic units (OTUs) were retained for analysis. To better compare community composition across samples we transformed the microbial count data to relative abundance. We analysed community composition at the family and genus levels for parent and offspring treatment groups using *phyloseq* (v1.46.0). α-Diversity was measured with the Shannon and Simpson indices (vegan v2.6.4), and dietary treatment effects were assessed with the Kruskal-Wallis test and Dunn’s pairwise comparisons (Table S2_AlphaDiversity). β-Diversity was evaluated using the Bray-Curtis dissimilarity matrix (abundance-based) via the *vegdist* function and hypothesis testing was conducted with PERMANOVA (Table S2_BetaDiversity). β-Diversity results were visualised with a non-metric multidimensional scaling (NMDS). The NMDS ordination was used to visualize the dissimilarity between samples, with MDS1 and MDS2 representing the main sources of variation in community composition. We fitted environmental factors (diet treatments) to the NMDS ordination, using the *envfit* function, and assessed the strength of the associations between bacterial taxa and treatment groups. To visualize these results, a heatmap was constructed, highlighting the most significant taxa that drive the observed patterns (p < 0.01, Table S2_BetaDiversity). We also performed a multilevel pattern analysis at the genus level to identify taxa associated with specific treatment groups (Table S2_Multilevel). Additionally, we generated a barplot depicting the family-level community composition, focusing on the top 20 taxa with the highest relative abundance.

To investigate potential relationships between gut microbial composition and gene expression in offspring, we selected a subset of taxa that contributed most significantly to differences among the dietary combinations in offspring (Figure 5b). A Spearman correlation analysis was then performed between these taxa and the most significantly differentially expressed genes identified in the gene expression analysis (Figure 4a and Table S2_LMM). We aimed to shed light on how parental diet might simultaneously shape the gut microbiota and gene regulatory networks in offspring, to explore potential communication pathways. The results were visualized with a heatmap using the *Hmisc* package (Figure 6).

## 3. Results

### 3.1 Intermittent fasting impacts on parental and offspring condition

The cross-mating combinations resulted in 18 successful families, this included five pregnant males in both AL(p)xAL(m) and IF(p)xIF(m) groups, and four for the AL(p)xIF(m) and IF(p)xAL(m) groups. In the remaining 14 cases, mating was unsuccessful due to either male rejection or early loss of eggs. Specifically, in the AL(p)xAL(m) group, two males discarded eggs within a day and one absorbed the few received; in IF(p)xIF(m), one male discarded the eggs and two refused to mate; in AL(p)xIF(m) two males absorbed eggs and two did not engage in mating; and in IF(p)xAL(m), one male absorbed the eggs and three did not mate. Morphological measurements, including length, weight, and fat content, were calculated for the successfully mated individuals, totaling 9 males and 9 females per treatment (AL or IF), and then weight and length for their offspring.

There was no significant difference in initial body length between any of the randomly assigned males or females at the start of the experiment (Figure S3a, Table S2_Morphology). Similarly, at the end of the experiment, final body lengths did not significantly differ between AL and IF groups within either sex (Figure S3a, Table S2_Morphology). However, when considering growth over the treatment period, both AL males and females showed greater increases in body length compared to their IF-treated counterparts. Using the median and standard error, AL males grew from 12 cm ± 0.44 to 13.5 cm ± 0.43 over the two-month diet period (P < 0.0001), and AL females from 12.6 cm ± 0.89 to 13.4 cm ± 0.79 over one month (P < 0.0001). In contrast, IF males showed a more modest increase from 12.5 cm ± 0.76 to 13.2 cm ± 0.78, and IF females from 12 cm ± 0.93 to 12.5 cm ± 0.87 (both P < 0.02) (Table S1 and S2_Morphology). A similar pattern was observed for body weight (Figure S3b). AL males increased from 0.62 g ± 0.07 to 0.94 g ± 0.09, while IF males showed a smaller change from 0.88 g ± 0.14 to 0.93 g ± 0.17. AL females grew from 0.84 g ± 0.24 to 0.89 g ± 0.19, whereas IF females showed a slight decline from 0.73 g ± 0.27 to 0.66 g ± 0.23. There was no significant difference between groups at the start of the experiment, but AL individuals showed a greater weight increase over the treatment period (P = 0.014), while IF individuals exhibited statistically insignificant weight changes (P > 0.8) (Table S1 and Table S2_Morphology).

In addition, we also assessed changes in overall body condition (Fultons Condition Factor). We found a significant overall effect of dietary treatment on the condition index at the end of the experiment (ANOVA, P < 0.002; Figure 1a). Post hoc comparisons revealed that only females from the AL and IF groups differed significantly (TukeyHSD, P < 0.03). Furthermore, linear mixed model analyses showed a significant decline in condition index over the fasting period for both IF females and males (P < 0.0001; Table S2_Morphology), reflecting the impact of food restriction. In contrast, AL individuals maintained a relatively stable condition index throughout the experimental period, despite some variation. For total body fat content in the parent pipefish, no significant treatment effect was observed between IF and AL females. However, there was a significant difference for males (P = 0.009, Figure 1b; Table S2_Morphology).

**Figure 1:**
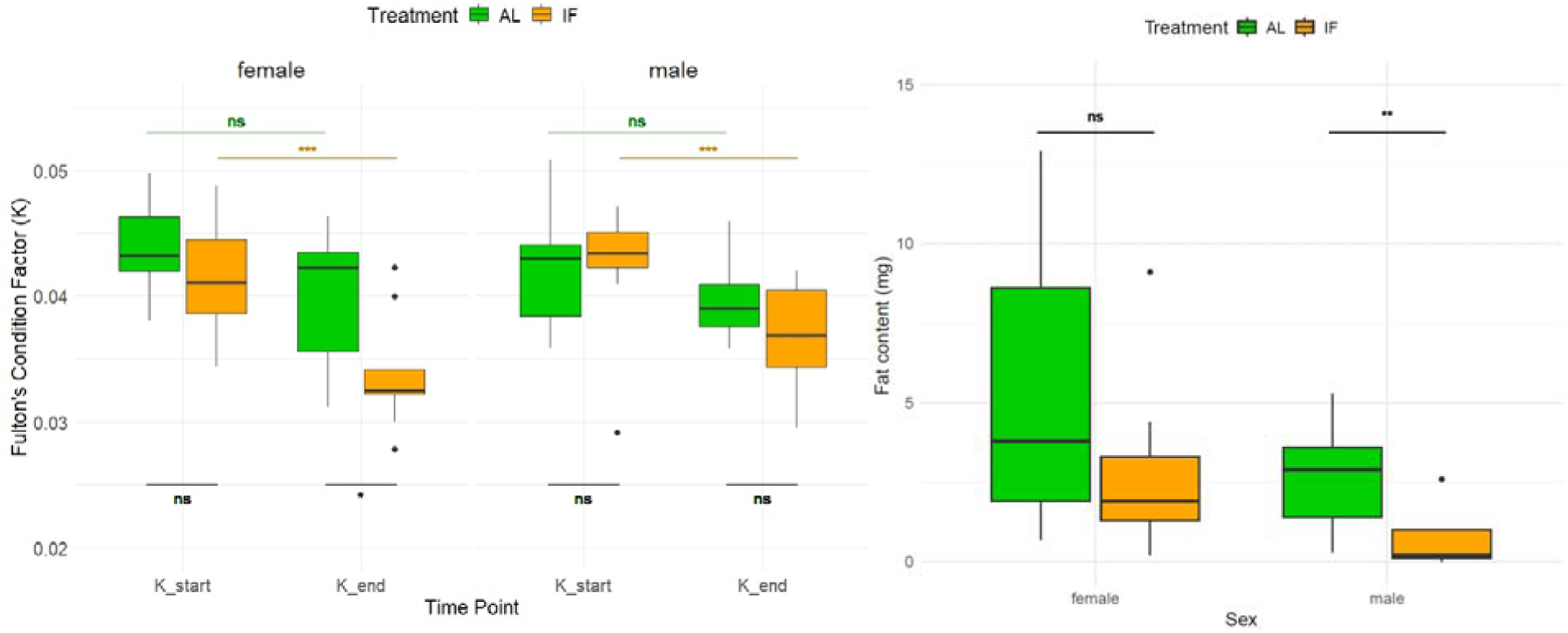
Condition index and fat content of *S. typhle* parents. a)The plot displays Fulton’s condition factor (weight/length³) for sex and dietary treatment, over time (x-axis). AL individuals are represented in green, and IF individuals are in orange. Two-way ANOVA results comparing treatment groups for both time points and sexes were not significant (ns). Significant differences are denoted as *p < 0.05, **p < 0.001, and ***p < 0.0001. b) Total body fat content (mg) of the pipefish parents. Females on the left had no significant difference in body fat content between AL and IF (P>0.1), while male pipefish were significantly different (P = 0.009, **).

A total of 5 juvenile per breeding pair were collected after the males gave birth, 20 each for the AL(p)xAL(m) and IF(p)xAL(m) group, 25 from the IF(p)xIF(m) group, and 16 juveniles for the AL(p)xIF(m) paring. The mean offspring length did not vary substantially across treatments, with AL(p)xAL(m) producing offspring with an average length of 20.97 ± 0.56 mm, IF(p)xIF(m) at 21.37 ± 0.55 mm, AL(p)xIF(m) at 23.21 ± 0.86 mm, and IF(p)xAL(m) at 21.84 ± 0.73 mm (P > 0.1; Table S1 and S2). Similarly, offspring weight was only marginally different between groups (P = 0.046), with AL(p)xAL(m) offspring averaging 5.49 ± 0.20 mg, IF(p)xIF(m) at 5.12 ± 0.18 mg, AL(p)xIF(m) at 5.83 ± 0.23 mg, and IF(p)xAL(m) at 5.89 ± 0.27 mg. After adjusting for multiple comparisons using Tukey’s HSD, none of the pairwise comparisons for weight were significant, with only the IF(p)xIF(m) showing a trend towards being lighter than IF(p)xAL(m) (P = 0.055, Table S2). When testing for Fulton’s factor we could not find a strong parental dietary effect on offspring body condition, and only a slightly lower K for offspring from AL(p)xIF(m) parents compared to AL(p)xAL(m) (Dunn test, P = 0.048, Table S2_morphology).

### 3.2 Intermittent fasting drives gene expression shifts in father’s and their offspring

To gain an overview of gene expression patterns across all samples and assess how variation was structured in the dataset, we first conducted a principal component analysis (PCA) using ΔCt values for both parents and offspring combined. This approach allowed us to evaluate whether treatment effects might be stronger than differences related to generation or sex. In the combined dataset, principal component 1 (PC1) explained 34.3% of the variance, primarily capturing the separation between offspring and parental samples, along with some plate effects (Figure S5a-b and Table S2_Permanova), while PC3 showed a marginal association with treatment (*P* = 0.058, Table S2). Given this clear generational distinction, subsequent analyses were conducted separately to disentangle treatment effects within each group. When focusing solely on offspring data, the association of PC3 with treatment effects strengthened (P = 0.019; Table S2_Permanova), indicating a clearer effect of parental dietary treatment on offspring gene expression. Post hoc pairwise comparisons (*adonis2*) showed a stronger difference between specific treatment groups, notably between IF(p)xIF(m) and AL(p)xIF(m) (P = 0.05), as well as IF(p)xIF(m) and AL(p)xAL(m) (P = 0.09) (Figure 4a and Table S2_Permanova).

Further analysis of individual gene expression responses to IF treatment in parents revealed varied effects in both sexes. In females, *DnMt3A* was the only gene with a significant response to fasting, reaching a log fold change (logFC) of 0.688 (LMM, P < 0.05; Table S2_LMM). In males, gene expression responses were more pronounced, particularly in immune-related pathways. Several markers of adaptive immunity—including *AIF*, *CD45*, *lymphocyt*, *IgM.lc*, and *TAP* were significantly upregulated in response to IF. Likewise, innate immunity genes *intf* and *tspo* also showed significant upregulation under IF treatment (Figure 3). Only one gene involved in both innate and adaptive immunity, *IL10*, was significantly upregulated (Figure 3). Among complement components, *C1* was the only gene downregulated by IF (Figure 3). Additionally, the regulatory gene *HDAC6* was also upregulated, pointing to a broader immunomodulatory effect of intermittent fasting in males.

In pipefish offspring several genes exhibited significant expression changes based on the parental treatment combinations, with AL(p)xAL(m) serving as the control. Within the adaptive immune response, *AIF* was significantly upregulated in the IF(p)xAL(m) group (logFC = 0.62, P = 0.05), while *bcell.rap31* was significantly downregulated in IF(p)xIF(m) offspring (logFC = –0.24, P = 0.03). For innate immune genes*, lectpt1* was strongly upregulated in the IF(p)xIF(m) group (logFC = 2.86, P = 0.03), and *lectpt2* showed substantial upregulation in both AL(p)xIF(m) (logFC = 5.36, P < 0.0002) and IF(p)xIF(m) offspring (logFC = 2.79, P = 0.03). The stress-related gene *hsp60* and *Intf* both showed non-significant trends toward downregulation in the IF(p)xAL(m) group (logFC = –0.39, P = 0.06; and logFC = –0.03, P = 0.06, respectively). Additionally, *ik.cytokine*, a gene implicated in both innate and adaptive immunity, was also downregulated in IF(p)xAL(m) offspring (logFC = –0.16, P = 0.057).

Regulatory genes were also significantly affected by parental diet treatments, particularly in offspring of IF-treated fathers (IF(p)xAL(m)). In this group, several regulatory genes were significantly downregulated, including *HDAC3* (logFC = –0.29, P = 0.03), *BROMO* (logFC = –0.26, P = 0.04), *JmjC_PHD* (logFC = –0.54, P = 0.02), *MYST1* (logFC = –0.49, P = 0.003), and *NO66* (logFC = –0.36, P < 0.0007). For both AL(p)xIF(m) and IF(p)xIF(m) group, *MYST1* was also downregulated (logFC = –0.39, P = 0.014; logFC = -0.36, logFC = 0.017), and *NO66* equivalently so (logFC = -0.26, P = 0.08; logFC = –0.34, P = 0.01, respectively). However, it is worth noting that not all of these genes met the ±0.5 logFC threshold (Table S2_LogFC), suggesting more moderate regulatory shifts in response to parental diet.

To further elucidate the influence of parental dietary combinations on offspring gene expression, we conducted an interaction analysis comparing -ΔCt values between AL and IF diets for both fathers and mothers for the most significant genes (Figure 4b-g).

When examining *Bcell.rap31* expression, the father’s diet played a dominant role: offspring exhibited low expression levels when the father was on an AL diet, regardless of the mother’s diet (Figure 4b). However, if the father was IF, *Bcell.rap31* expression in offspring increased significantly, suggesting a paternal-driven effect on this adaptive immune gene. Also *Hsp.60* demonstrated a more paternal-driven influence, with expression levels consistently lower when the father was fed AL, irrespective of the mother’s diet (Figure 4f). Offspring expression of *Hsp.60* increased only when the father was on an IF diet, underscoring the father’s dietary intake as the main driver.

Patterns in the innate immune genes *Lectpt1* and *Lectpt2* mirrored each other closely (Figure 4c and Figure S5a). Expression was highest in offspring when both parents were on an AL diet, while offspring from an IF mother with an AL father showed the lowest levels. Interestingly, mixed diet pairs (*AL(p)xIF(m)* and *IF(p)xAL(m)*) resulted in intermediate expression levels, suggesting that parental diet mismatch modulates gene expression toward a balance between the extremes.

For *Intf*, the maternal diet emerged as the primary influencer: offspring exhibited higher expression when the mother was on an AL diet, with expression levels dropping under an IF maternal diet (Figure 4e). This trend persisted regardless of the father’s diet, indicating a maternal dominance in regulating *Intf* expression.

In the case of *Ik.cytokine*, parental IF diets synergistically amplified expression, with the highest levels observed when both parents were on IF (Figure S5b). Even when only one parent was on an IF diet, *Ik.cytokine* expression increased, suggesting that any exposure to IF in either parent potentiated this gene’s expression, with dual-IF parental treatment yielding the greatest effect.

A similar interaction was observed for *AIF*, which showed elevated expression when both parents shared the same diet, either AL or IF (Figure 4c). However, expression of *AIF* decreased when parental diets differed, hinting at a synchrony effect where parental dietary alignment might maximize expression levels.

Interestingly, the regulatory genes followed a consistent pattern, irrespective of their specific roles in gene activation (*BROMO* and *MYST1*) or gene silencing (*NO66*, *JmjC-PHD*, and *HDAC3*). Across all regulatory genes, offspring from AL(p)xAL(m) parents showed the lowest expression levels, while those from an IF father and AL mother consistently exhibited the highest (Figure S5c-f). The mother’s fasting status appeared to have minimal impact, causing only slight fluctuations when she was IF, suggesting a significant paternal influence on regulatory gene expression in offspring.

### 3.2 Fasting affects microbial composition of father’s and offspring

Microbial alpha diversity metrics revealed no significant differences across the groups - neither between parents and offspring nor between treatment groups (AL vs. IF) (Table S2_AlphaDiversity). This suggests that within-group microbial richness and evenness remained stable. In contrast, beta diversity analysis showed significant differences across groups (PERMANOVA, P < 0.001). Pairwise comparisons revealed that the strongest differences occurred between parents and offspring (Table S2_BetaDiversity), as illustrated in the NMDS plot (Figure 5), where NMDS1 clearly separates the generational differences.

Further inspection within the adult groups showed no significant differences in beta diversity between AL males and AL females or between AL females and IF females (Table S2_BetaDiversity). However, a significant difference emerged between AL males and IF males (P.adj = 0.013), indicating that fasting influenced microbial community composition in male pipefish but not in females. This distinction is also reflected in Figure 5, where IF males tend to cluster toward the upper region, while other adults are generally concentrated toward the bottom left. Among offspring groups, the most pronounced differences in microbial composition were between the AL(p)xIF(m) and other offspring pairings (P.adj < 0.05).

The bacterial genera driving the observed separation between parental and offspring gut microbiomes are illustrated in Figure 5b (Table S2_BetaDiversity). Notably, certain genera - such as *Vicingus* (*Cryophormacea*), *Spongiivirga* (*Flavobacteriaceae*) and *Sulfitobacter* (*Rhodobacteraceae*) - were consistently more abundant in parents compared to offspring (Figure 5b), reinforcing the distinct microbial profiles identified by beta diversity analysis. For instance, *Vicingus* was present at higher relative abundances in adult samples (7-15%) but remained below 0.4% in offspring samples (Figure 5c). Conversely, *Kordia*, *Pseudoalteromonas, Aliikangiella,* showed greater relative abundance in offspring, with *Pseudoalteromonas* appearing more prominently in offspring of fasting parents (Figure 5 b & c).

Within the parental groups, diet also shaped microbial composition, as evidenced by increased *Brevinema* in AL fed parents and higher *Pseudoalteromonas* in IF parents (Figure 5c). Among offspring, combinations of different parental diets (AL(p)xIF(m) or IF(p)xAL(m)) and similar ones clustered together (Figure 5a&b), with example decreased relative abundances of *Dokdonia, Kiloniella*, and *Porticoccus* for IF(p)xAL(m) and AL(p)xIF(m).

The correlation analysis between the genera *Brevinema, Marinomonas, Porticoccus, Kiloniella, Dokdonia*, and the significantly differentially expressed genes yielded only a few significant associations (Figure 6). The strength and direction of these associations varied across dietary combinations, with more significant patterns observed in AL(p)xAL(m) and IF(p)xIF(m) offspring compared to the mixed diets (Figure 6). Offspring from the AL(p)xAL(m) parental dietary combination exhibited strong correlations between specific microbial taxa and immune-related gene expression. *Brevinema* showed a negative correlation with *Ik.cytokine*, while *Dokdonia* was positively correlated with *AIF* and *Intf*. Additionally, *Porticoccus* demonstrated a strong positive correlation (>0.8) with *Lectpt1* and *Lectpt2*. Although these associations were weaker and not statistically significant in the IF(p)xIF(m) group, the trend remained positive. In mixed dietary groups (AL(p)xIF(m) and IF(p)xAL(m)), *Porticoccus* exhibited negative correlations with *Lectpt1* and *Lectpt2*, though these associations were also not significant. *Dokdonia* emerged as a recurring key player, with positive correlations observed with *Intf* in both AL(p)xAL(m) and IF(p)xIF(m) offspring and with *NO66* in IF(p)xAL(m). *Marinomonas* showed contrasting trends, being positively correlated with *Hsp.60* in AL(p)xIF(m) but negatively correlated with the same gene in IF(p)xIF(m).

## 4. Discussion

Shifting environmental conditions frequently require organisms to rely not only on immediate plastic responses, but also on TGP where survival often depends on the intergenerational transfer of biological experience (1). This inherited knowledge, encoded within physiological, epigenetic marks, microbial communities, or behavioural cues, can improve offspring-environmental matching and potentially serve to evolutionary rescue (69). The current understanding of how parents influence their offspring is predominantly based on conventional sex roles, where mothers bear the responsibility of both the costly gametes and pregnancy. However, nature evolved a diverse array of reproductive strategies and parenting roles. Reproductive strategies might profoundly influence non-genetic inheritance, i.e., parental care might enhance the opportunity for sex-specific trans-generational transfer fostered by intimate contact of one parent to the offspring. The timepoint of transfer of parental experience, e.g., whether during the fusion of egg and sperm, or, alternatively, during pre- or postnatal parental care, transfer of parental experience occurs, might additionally shape the outcome and role of non-genetic inheritance in offspring life. To gain a comprehensive understanding of how parental experiences shape subsequent generations, we must expand our perspective beyond the familiar and explore systems where the functions of sex, care, and gestation are dissociated.

The broadnosed pipefish, with its reversed sex roles and male pregnancy, presents a rare opportunity to disentangle the effects of maternal provisioning from the physiological impacts of gestation (48,57,62). This species’ exceptional reproductive strategy makes it an ideal model for exploring how sex-specific parental environmental stressors shape offspring phenotypes (48,61,70). By leveraging this distinct reproductive system, our study aimed to disentangle the relative contributions of maternal provisioning and the paternal gestational environment to offspring development, offering novel insights into how sex-specific and gestational effects interact under nutritional stress. In the present study, we examined the influence of parental diet on offspring development by exposing both male and female *S. typhle* to either intermittent fasting (IF) or ad libitum (AL) feeding prior to mating, and continuing the IF regime during male pregnancy. We then assessed morphological traits, immune and epigenetic gene expression, and gut microbiota composition in both parents and their offspring. While our design captures both maternal and paternal contributions, including paternal care via male pregnancy, it is possible that some of the observed offspring effects stem from early, pre-fertilization influences such as epigenetic modifications in gametes. Distinguishing between such pre- and post-fertilization mechanisms is a key future goal which requires comparative experimental approaches between closely related species along a parental care gradient. This study provides a first stepping stone toward disentangling the relative roles of gametic programming and parental provisioning in shaping offspring phenotype.

The IF regime imposed a clear energetic cost, as evidenced by reduced growth in length and weight (Figure S3a&b), and declines in Fulton’s condition index (K) in both sexes over time (Figure 1a). Notably, only IF females were in a weaker endpoint condition compared to their AL counterparts (P < 0.03), despite males also exhibiting fat loss (P = 0.009, Figure 1b) and suppressed growth (Figure 1a). This is particularly striking given that males were subjected to the IF regime for twice as long as females. Females showed no reduction in fat content (Figure 1b), suggesting that the decline in K may reflect loss of lean tissue such as muscle. This female-specific sensitivity is striking, particularly given that males where fasted for twice as long. A similar pattern has been reported in *Hippocampus erectus*, where caloric restriction affected condition in females but not males, and was linked to female-biased metabolism and fat storage strategies (56). Such sex-specific responses are conserved across taxa: in mice, for instance, males tend to preserve muscle and bone by mobilizing fat stores, while females resist fat loss and instead catabolize lean mass, often alongside reproductive suppression (71,72). Our findings suggest that female pipefish may similarly be more vulnerable to fasting, possibly due to sex-specific energy allocation strategies. Alternatively, males may have compensated for long-term restriction through physiological or behavioural adjustments.

Although offspring morphology was only marginally affected by parental dietary treatment - mean lengths across groups were fairly similar (∼21–23 mm), and IF(p)xIF(m) offspring trended towards lower weight - we detected a significant difference in offspring condition (Figure 2). Specifically, offspring from AL(p)xIF(m) crosses had a lower condition index compared to those from AL(p)xAL(m) crosses (Dunn test, P = 0.048; Figure 2). Since juveniles were measured immediately after birth, it is possible that latent or delayed morphological effects could emerge later in development (e.g., in growth rate, behaviour, or survival). Nonetheless, these findings indicate that maternal physiological state or egg provisioning plays a crucial role in shaping early juvenile condition, even in a species that exclusively relies on post-fertilization paternal care. A well-fed father (AL) was unable to fully buffer against the consequences of reduced maternal input (IF), while offspring from fasted fathers and well-fed mothers (IF(p)xAL(m)) showed no such deficit, even though paternal feeding conditions remained throughout pregnancy. This asymmetry highlights the central role of maternal investment in early development in fish (73,74). While males in *S. typhle* contribute significantly through gestation, egg quality remains a limiting factor (56,70). Our results suggest that maternal provisioning can more effectively compensate for suboptimal paternal condition than vice versa, possibly because females are able to prioritize and sustain ovarian function even under IF - drawing on somatic reserves at the expense of body condition -while incurring a lower energetic cost than males investing in pregnancy.

**Figure 2:**
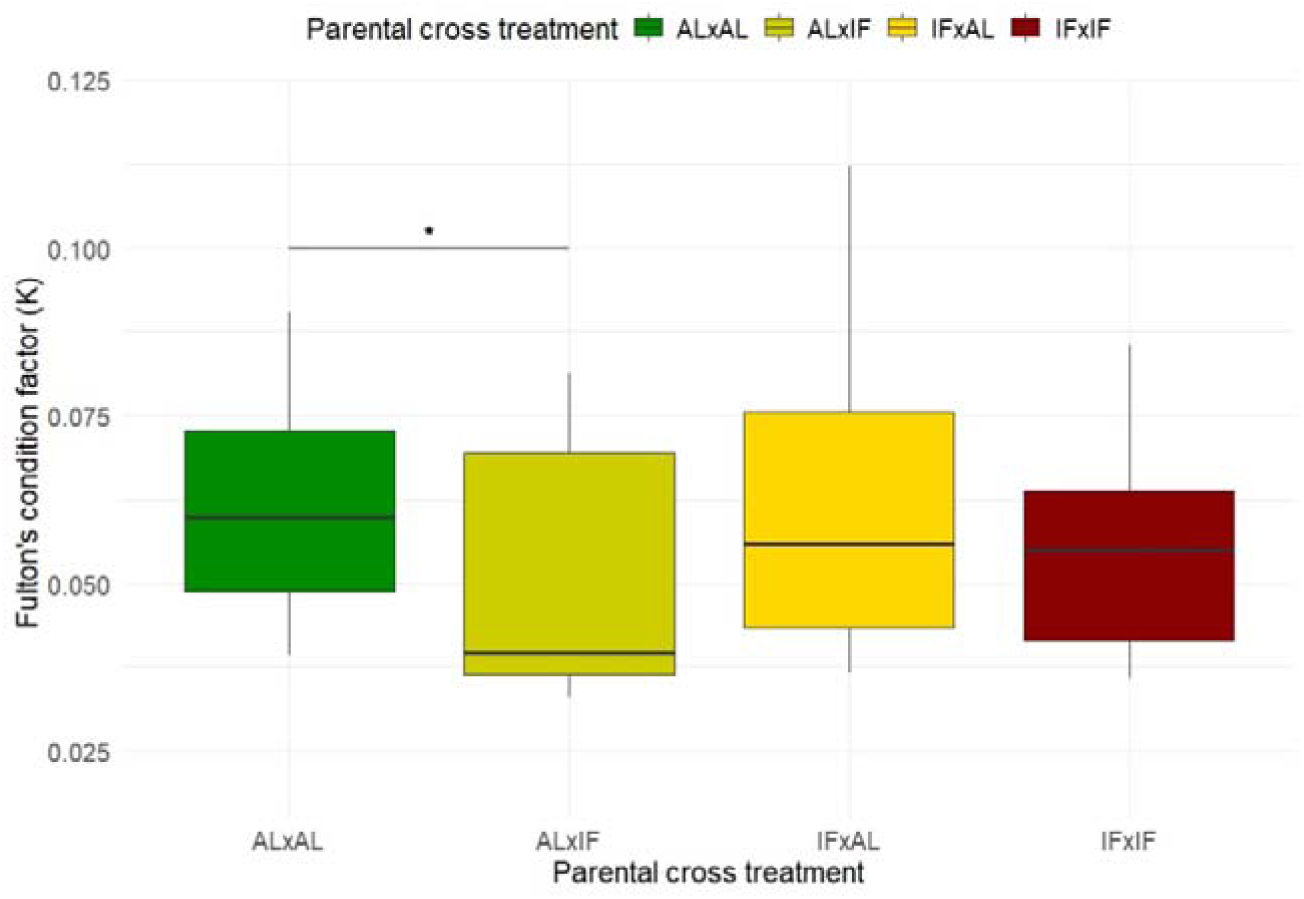
Offspring physiological condition at birth. Plot illustrates Fulton’s condition factor for offspring at birth across different parental dietary treatment groups: dark green for AL×AL, red for IF×IF, dark yellow for AL×IF, and yellow for IF×AL.

Principal component analysis of gene expression profiles showed that variation was predominantly structured by life stage (offspring vs. parents), with only subtle effects of dietary treatment emerging on PC3 (Figure S4a-c and Figure 4a). In females, *DnMt3A* - a key regulator of *de novo* DNA methylation (75)-was the only gene to surpass both the significance threshold and a logFC of 0.5 under IF treatment. All adult fish were sampled immediately after completing their respective reproductive investment (i.e., egg production in females, pregnancy in males) to capture gene expression signatures at the biologically relevant endpoint of parental allocation. The modest transcriptional response in females may reflect a combination of shorter IF exposure and sex-specific physiological strategies. Indeed, this limited change aligns with our phenotypic observations, where females exhibited reduced condition but no measurable fat loss, suggesting distinct energy mobilization and possibly constrained regulatory flexibility under dietary stress. In contrast, IF males exhibited a broader molecular shift with pronounced upregulation of genes associated with the adaptive immune response, including *AIF, CD45, TAP*, and *IgM.1c*, alongside elevated expression of anti-inflammatory cytokines (*IL10* and *intf*) and a downregulation of the complement component *C1* (Figure 3). This transcriptional profile is consistent with established immunological responses to intermittent fasting. IF is known to enhance autophagy, a cellular process that facilitates the clearance of intracellular pathogens and damaged components, while reducing oxidative stress through decreased ROS levels and increased SIRT1 expression (76,77). Furthermore, the observed downregulation of *C1*, a component of the classical complement pathway, aligns with the shift toward an anti-inflammatory and possibly more immune-tolerant state under energetic constraint. The upregulation of CD45 - a pan-leukocyte marker critical for T and B cell receptor signalling - further suggests that fasting stimulates immune cell viability and activity, possibly through autophagy-mediated pathways. In mammals, short-term intensive fasting has been shown to enhance the survival and function of CD45+ leukocytes by reducing apoptosis and promoting cytokine secretion (76,78), a mechanism that may be evolutionarily conserved and similarly activated in *S. typhle*. These findings suggest that male pipefish adjust their immune expression more readily than females under nutritional stress, perhaps reflecting the physiological demands of male pregnancy. Under nutritional stress, this balance might shift -forcing the immune system to compensate more strongly to maintain both pouch health and self-protection. Reduced investment in growth or fat stores under IF allows reallocation of limited resources to immune defence, especially in pregnant males.

**Figure 3.**
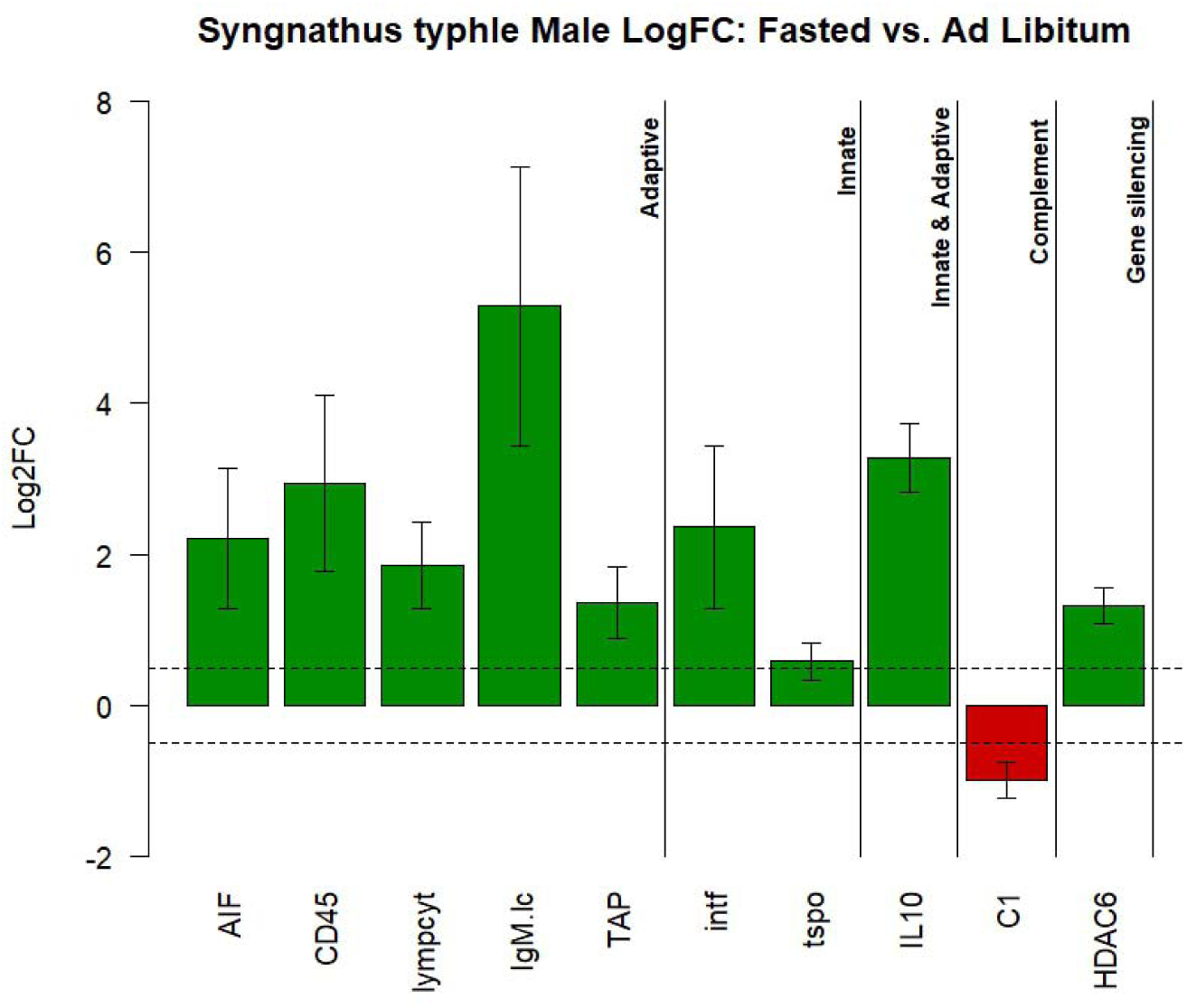
Bar plot of LogFC values for male *Syngnathus typhle* fathers, comparing IF to AL feeding as a control. The bar plot displays only those genes that were significant in the linear mixed model (LMM) analysis (Table S2) and met the LogFC threshold of ± 0.5. Genes upregulated in fasted males are shown in green, while downregulated genes are in red. The genes are categorized into functional groups, including immune system categories (adaptive, innate, innate & adaptive, complement) and gene regulatory groups (e.g., gene silencing). Of the 10 genes depicted, 9 are upregulated in fasted males relative to controls, with C1 being the sole gene downregulated.

The upregulation of immune-related genes in IF males coincided with significant shifts in gut microbial composition, suggesting a potential link between microbial signals and host immune modulation. Specifically, IF males showed increased abundance of *Pseudoalteromonas*, a genus within Proteobacteria also found to dominate in calorie-restricted seahorses (56). *Proteobacteria* are known to support gut health by promoting epithelial integrity and facilitating colonization by strict anaerobes, while also influencing mucosal immune responses (79). In contrast, AL-fed males displayed higher levels of *Brevinema*, a genus previously associated with fish mucus (80) and notably implicated in paternal-specific microbial transmission in Syngnathids (15). This microbial shift may reflect not only differences in dietary input, but also immunological state, as *Brevinema* dominance in AL males was paralleled by reduced expression of immune genes relative to the more activated immune profile of IF males. Given that IF has been shown to enhance epithelial barrier function and modulate immune responses through microbiota-driven mechanisms, it is plausible that the microbial changes observed here contribute to the transcriptional immune shifts in IF males (76,81). Moreover, the reduction in *Brevinema* under IF may indicate a disruption in paternal microbial transfer potential, possibly linked to altered immune signalling or reduced capacity for microbial hosting during reproduction.

Gene expression analyses in pipefish offspring revealed nuanced yet meaningful immunological and regulatory responses to parental dietary history. These effects were not uniform across gene types or treatment combinations, but instead appeared to reflect parent-specific transmission of dietary stress signals, with IF(p)xAL(m) and IF(p)xIF(m) offspring showing the strongest expression shifts. The consistent upregulation of chromatin remodelling and transcriptional regulatory genes (e.g., *HDAC3, JmjC-PHD, MYST1, BROMO, NO66*; Figures S5c-g) in offspring of IF-fed males suggests that paternal nutritional stress may prime offspring for broader developmental plasticity or environmental responsiveness, perhaps via epigenetic reprogramming. This is consistent with findings in other taxa where paternal diet has been shown to influence offspring metabolism, behaviour, and disease susceptibility through modifications such as DNA methylation and histone acetylation (35,82,83). Although genome-wide epigenetic reprogramming occurs post-fertilization, certain epigenetic marks can be maintained across generations - particularly at imprinted loci or other regulatory hotspots - allowing paternal nutritional experiences to influence early developmental trajectories (82). This could enable the transmission of anticipatory cues to offspring, potentially shaping their metabolic or immune responsiveness in environments predicted to be resource-limited. Additionally, both *Bcell.rap31* – essential for B cell development and proliferation (84) – and *Hsp60 -* a mitochondrial stress-related gene (85) - showed increased expression in offspring when fathers were IF, regardless of maternal diet (Figures 4b and f). *Hsp60* functions primarily as a mitochondrial chaperone but also contributes to broader cellular processes including cell proliferation, apoptosis, migration, and immune regulation (86). The simultaneous upregulation of these genes suggests that paternal fasting may act as a cue, priming offspring for anticipated physiological stress by enhancing immune readiness. This points to a form of intergenerational plasticity, where paternal nutritional status modulates offspring physiology through inherited immunological components (9,62).

**Figure 4:**
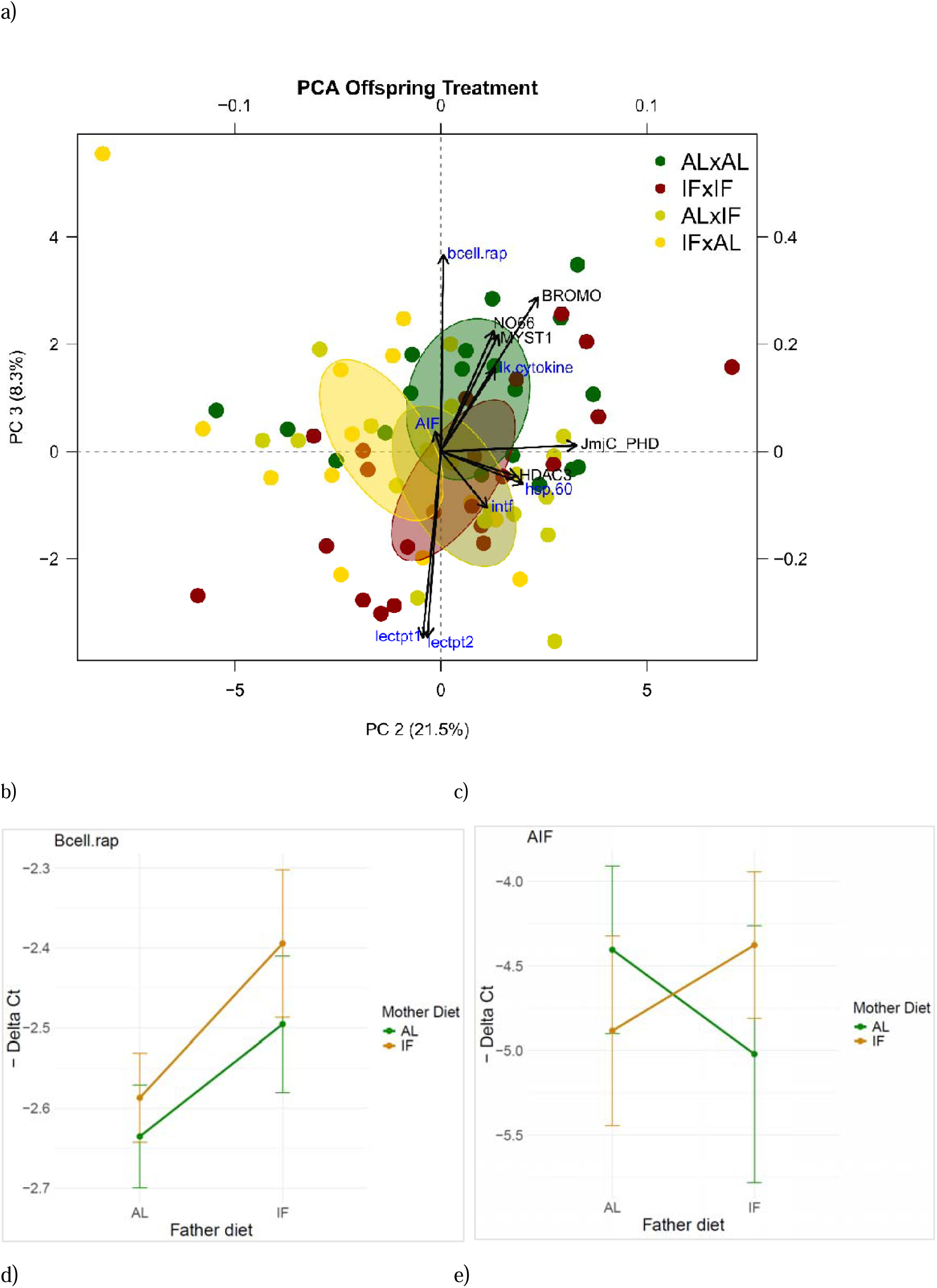
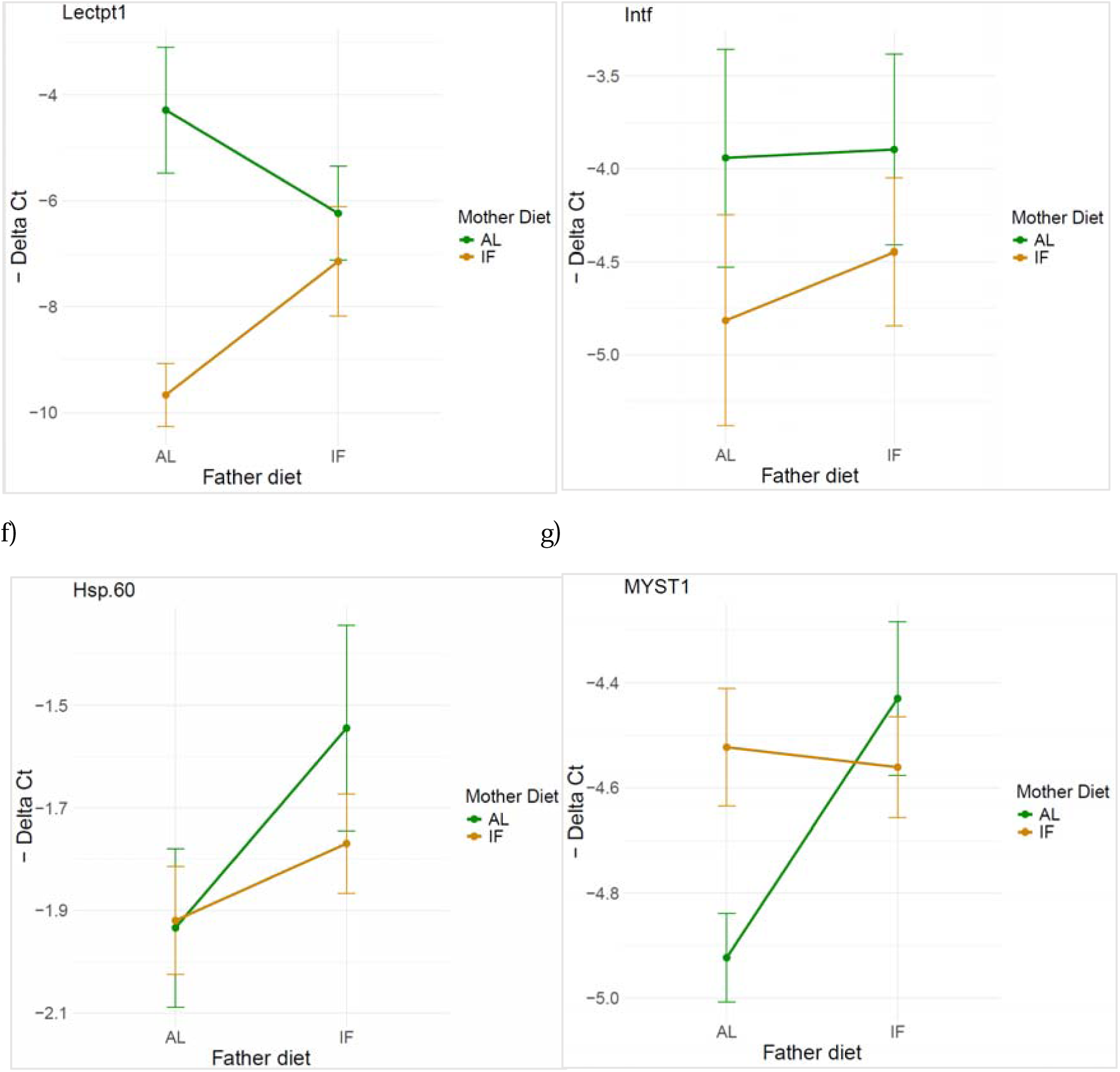
Parental dietary treatment effects on offspring gene expression. a) PCA plot of differentially expressed genes in offspring, based on parental dietary combinations. The x-axis and y-axis represent PC2 and PC3, explaining 21.5% and 8.3% of the variance, respectively. Secondary axes display the gene loading coordinates, with arrow lengths scaled to reflect each gene’s contribution to these principal components. This plot highlights only those genes identified as differentially expressed through a linear mixed model (LMM) analysis (see Table S2_LMM). Genes categorized as regulatory are labelled in black, while immunological genes are in dark blue (b) to (g) show interaction plots for significantly differentially expressed genes in *S. typhle* offspring, including Bcell.rap31 (b), AIF (c), Lectpt1 (d), Intf (e), Hsp.60 (f), and MYST1 (g). Additional genes, including Lectpt2, Ik.cytokine, BROMO, HDAC3, NO66, and JmjC-PHD, are detailed in the supplementary material (Figure S5a-f). In these plots, the x-axis represents the father’s diet, while colour coding (green for ad libitum and orange for intermittent fasting) indicates the mother’s diet. The y-axis displays the negative Delta Ct values, reflecting the directionality of gene expression. Each plot illustrates the interactions between parental dietary treatments and their effects on offspring gene expression.

Rather than enhancing offspring immune preparedness, maternal IF appeared to dampen innate immune gene expression in the next generation. Offspring of IF-fed mothers showed consistent downregulation of key innate immune genes such as *lectpt1, lectpt2*, and *intf* (Figures 4d, S5a–b), regardless of paternal diet. Additionally, *ik.cytokine* was upregulated in these same offspring (Figure S5b), a gene associated with the suppression of IFN-gamma, potentially through downregulation of MHC class I expression (87). This pattern suggests that maternal fasting may constrain investment in early immune programming, possibly due to energetic limitations during oogenesis. This interpretation is supported by other findings in *S. typhle* showing that environmental stressors like elevated temperatures can erase transgenerational immune priming, likely reflecting trade-offs between survival and immune investment when resources are limited (48,57). Our results may reflect a similar mechanism, where maternal stress cues lead to reduced immune provisioning as an adaptive reallocation of resources under caloric constraint.

Parental dietary treatment not only shaped offspring gene expression but also influenced their gut microbial composition. Taxonomically, offspring microbiomes differed markedly from those of adults, with genera like *Vicingus*, *Spongiivirga*, and *Sulfitobacter* (taxa associated with marine environments (88,89)) abundant in parents but nearly absent in offspring (Figures 5b-c). It is crucial to highlight that these differences may not solely reflect parental dietary influences, juveniles naturally consume a very different diet at birth (e.g., copepods), whereas adults feed on larger invertebrates (90). As such, developmental dietary divergence likely contributes to these microbial shifts. Conversely, *Pseudoalteromonas*, *Kordia*, and *Aliikangiella* were more prevalent in juveniles, with *Pseudoalteromonas* particularly enriched in offspring of fasting parents, mirroring IF parents and suggesting a microbial legacy of parental nutritional state (Figures 5b-c). This genus includes marine species known for a wide range of functions, such as antibacterial, algicidal, and antiviral activities, as well as specific strains that prevent the settlement of common fouling organisms (91).

**Figure 5:**
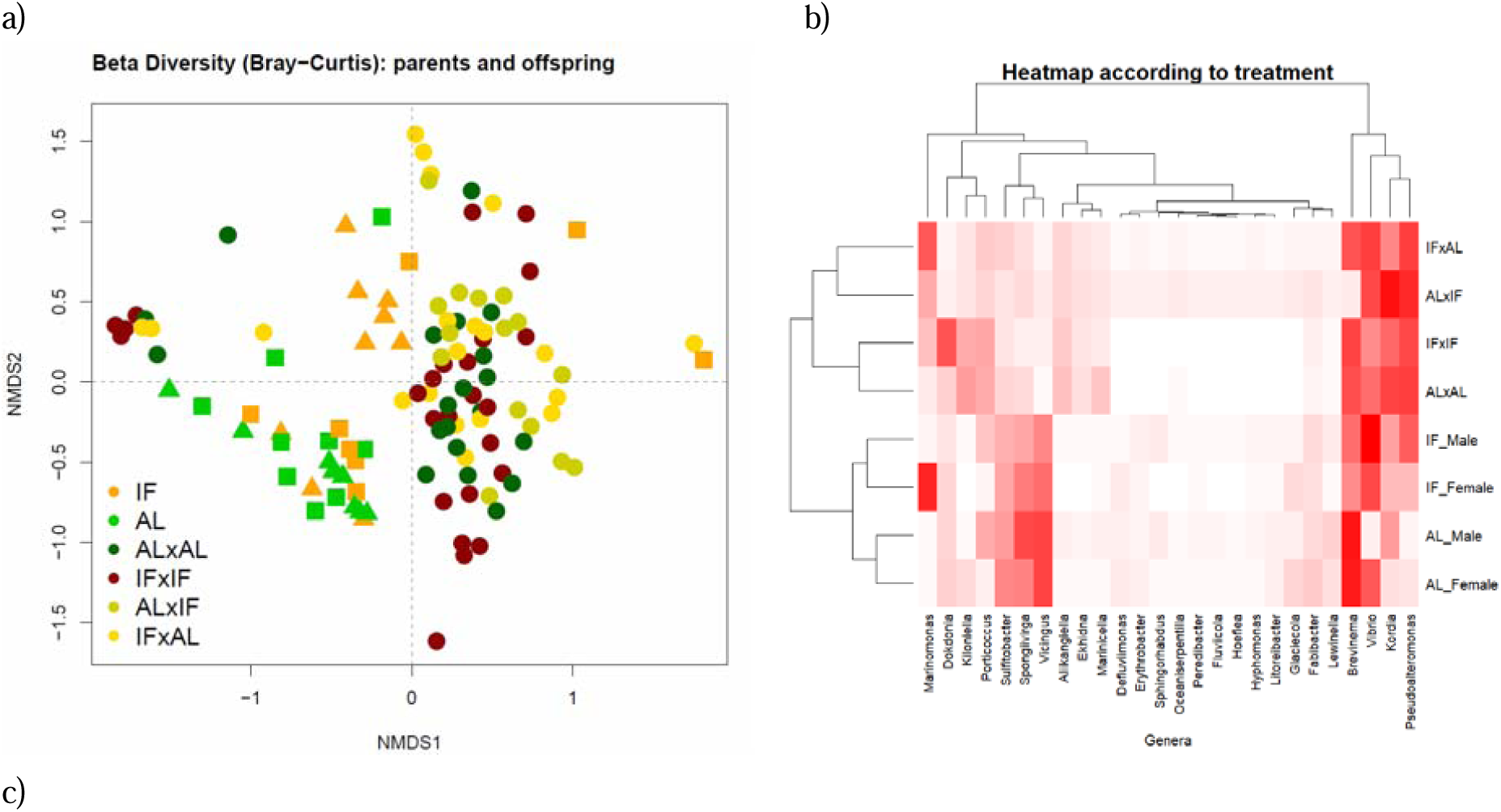
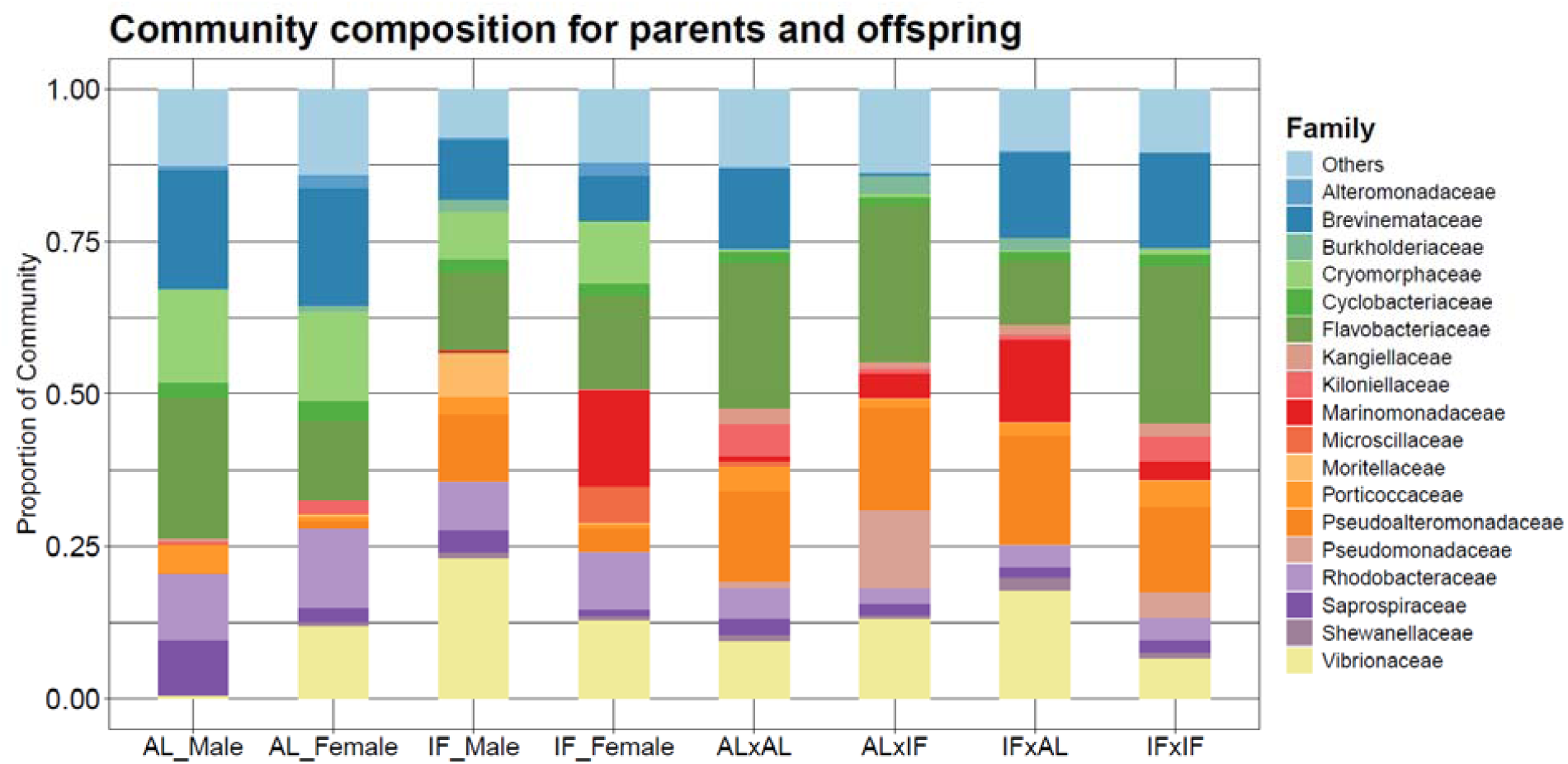
a) Bray-Curtis NMDS plot illustrating bacterial community composition in pipefish parents and their offspring. Points are coloured by diet: fasting parents are shown in orange, and ad libitum parents in light green. Offspring color-coding reflects parental diet combinations, with dark green for offspring from both AL parents, dark red for offspring from both fasting parents, olive green for offspring of an AL male and IF female, and yellow for offspring of IF male and AL female. Parental sex is denoted by symbols: an upward triangle represents males, and a square indicates females. NMDS1 shows clear separation between offspring and parents, highlighting distinct bacterial community compositions across generations. b) Heatmap of the most significant (P < 0.01) taxa driving Bray-Curtis beta diversity patterns. Taxa abundance is shown by red intensity, with darker shades indicating higher relative abundance. The y-axis dendrogram clusters samples based on treatment groups, while the x-axis dendrogram clusters taxa by similarity. c) Bar plots showing the proportional composition of bacterial communities at the family level, highlighting the 20 most abundant taxa across all groups.

Nonetheless, PCA and beta diversity analyses revealed that offspring microbiomes clustered by parental diet combinations (Figure 5a–b). Offspring from mixed-diet crosses (AL×IF, IF×AL) harboured increased levels of *Marinomonas*, whereas those from matched parental diets (AL×AL and IF×IF) showed higher relative abundances of *Dokdonia, Kiloniella*, and *Porticoccus*. Correlation analysis (Figure 6) further indicated that *Marinomonas* was positively associated with the stress-responsive gene *Hsp60* only in AL×IF offspring, but negatively associated in both matched-diet groups. Meanwhile, *Porticoccus* was strongly positively correlated with innate immune genes *lectp1* and *lectp2* in AL×AL offspring, and similarly (though not significantly) in IF×IF, but not in offspring from mixed-diet crosses. Likewise, *Dokdonia* showed positive associations with immune markers *AIF* and *intf* in IF×IF, AL×AL, and (non-significantly) AL(p)xIF(m) offspring, but this association reversed in IF(p)xAL(m) offspring – where the father experienced IF. This reversal could imply that paternal fasting, when not mirrored by the mother, disrupts coordinated immune-microbiome interactions, potentially altering offspring gut environment. In a species like *S. typhle*, where both mothers and brooding fathers contribute to the microbial inoculum - via eggs and brood pouch environments - matched parental diets may support the maintenance or co-transfer of a similar microbial community (49,92). Furthermore, when diets match, parental epigenetic and physiological cues (e.g., immune tolerance, gut pH/mucosal environment, antimicrobial peptides) may be aligned, producing a gut environment in the offspring that selects for a consistent microbial community. With mismatched diets, the offspring may receive conflicting cues - say, maternal tolerance toward one microbial profile and paternal priming toward another - leading to disrupted or restructured microbiome assembly.

**Figure 6:**
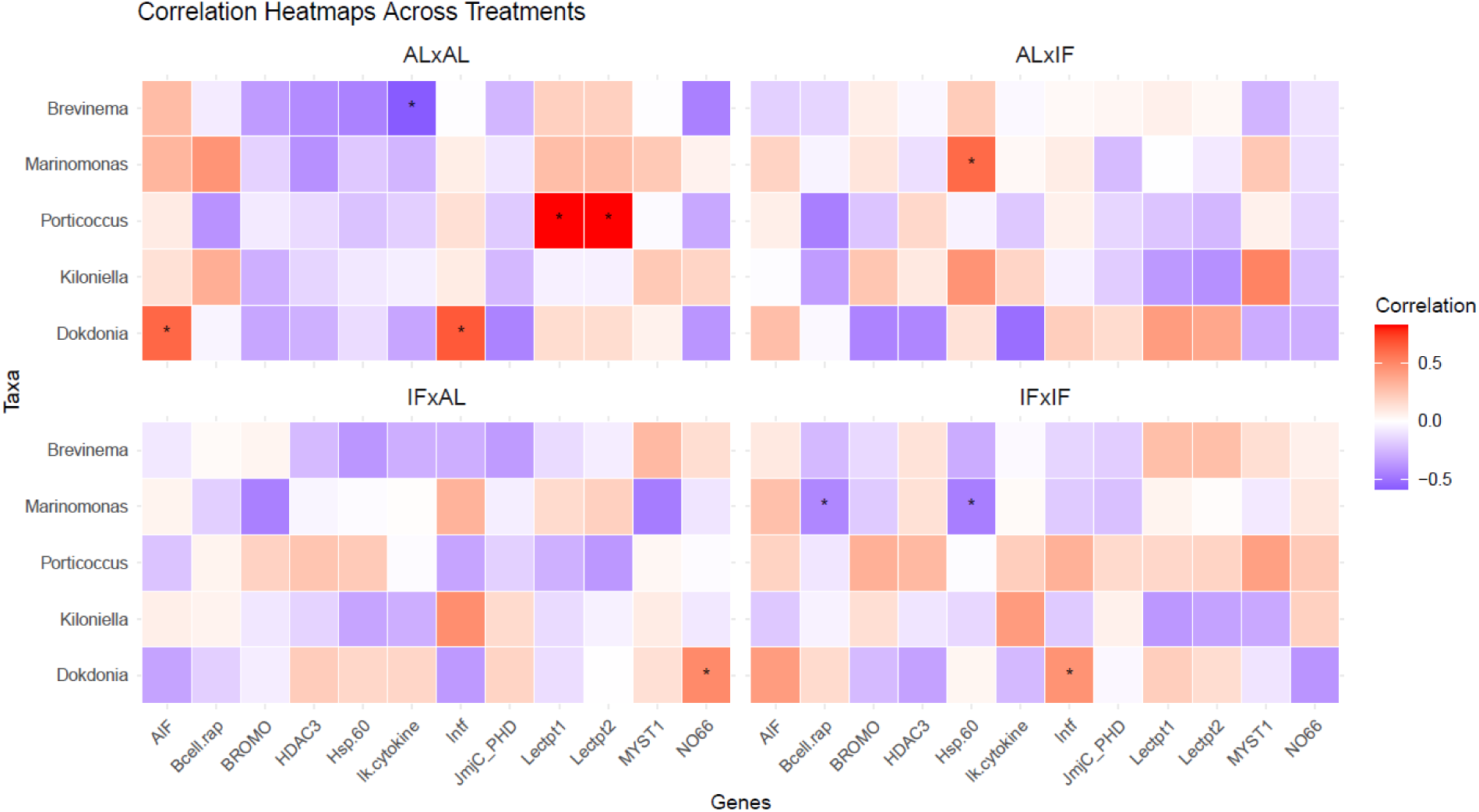
Spearman correlation heatmap depicting the relationships between significant differentially expressed genes and microbial taxa likely driving offspring treatment divergence, for offspring from the four different parental dietary treatment groups. The x-axis represents the genes, while the y-axis displays the microbial genera. Colour intensity reflects the strength of the correlation, with dark blue indicating r < -0.5 and dark red indicating r > 0.5. Asterisks (*) highlight statistically significant correlations (p < 0.05), emphasizing key relationships between gene expression and microbial composition in the context of parental diet treatments.

## Conclusion

Our study revealed how parental nutritional stress shapes offspring phenotype through sex-specific and asymmetric pathways in the sex-role reversed pipefish *Syngnathus typhle*. By manipulating maternal and paternal diets, we demonstrate that maternal provisioning plays a dominant role in determining offspring condition, while paternal effects emerge more clearly in immune gene expression and microbiota composition. This highlights the importance of considering both maternal and paternal contributions—especially in systems where traditional sex roles are decoupled.

Notably, females experienced greater decline in condition index under IF, which translated into lower offspring condition, regardless of paternal diet. In contrast, offspring of IF males and well-fed females showed no such deficit, underlining the critical role of maternal provisioning in early development. However, IF fathers did not remain passive contributors; they exhibited strong immune activation and altered microbiomes, both of which were reflected in offspring gene expression and microbial profiles. These findings suggest that paternal nutritional stress can prime offspring immune function, likely through epigenetic and microbial inheritance mechanisms. Importantly, mismatched parental diets disrupted expected immune-microbiome correlations in offspring, indicating that coordinated parental cues may be essential for stable microbial and physiological development. Such interactions between diet, immunity, and microbiota underscore the ecological relevance of parental environmental history in shaping offspring resilience.

This research leverages the unique reproductive biology of *Syngnathus typhle* to disentangle the often-confounded effects of maternal provisioning and gestational environment under nutritional stress. It contributes a unique perspective to the study of intergenerational plasticity by moving beyond conventional reproductive models. Understanding these dynamics is critical in an era of rapid ecological change, where nutritional stress and altered resource availability are increasingly common through climate change. *S. typhle* provides a powerful model for investigating how sex-specific and gestational pathways interact to mediate offspring outcomes across generations.

## Acknowledgements

Special thanks goes to Silke-Mareike Marten and Diana Gill for their valuable input in the laboratory. We are also very grateful to Dr. Ralf Schneider, who provided useful insights and feedback on the Fluidigm and statistical analysis. Financial support was provided by the Deutsche Forschungsgemeinschaft (DFG, German Research Foundation) through the Research Training Group for Translational Evolutionary Research (RTG 2501 TransEvo), and through the SynSex project RO 4628/1-8 to OR, by the European Research Council (ERC) under the European Uniońs Horizon research and innovation program (MALEPREG: eu-repo/grantAgreement/EC/H2020/755659) to OR, and through the CRC 1182 Origin and Function of Metaorganismus for supporting microbiota genotyping.

## Data availability statement

More information can be found in the supplementary material, including information on methodology and additional figures (Figures S1-S5). Table S1 contains morphological raw data on length, weight, fat content and condition index of parents and offspring of *S. typhle*. Tables S2 displays the full statistical results in several panels for morphology, gene expression analysis, permanova analysis, the logFC results, results for the alpha and beta diversity analysis. Table S3 contains information on the primers we used for the Fluidigm analysis and their efficiency values. Table S4 has the raw CT values from the Fluidigm analysis. Table S5 has the raw genus level count data, metadata and tax information. The raw sequencing 16S rRNA-Seq and metadata used in this study are available from the National Center for Biotechnology Information (NCBI) Sequence Read Archive (SRA) under BioProject ID PRJNA1270975 - SUB15360138.

## Ethics statement

All experimental procedures complied with German animal welfare regulations and were approved by the Schleswig-Holstein authority Ministerium für Energiewende, Landwirtschaft, Umwelt, Natur und Digitalisierung (MELUND) under permit number 1368/2023. The study did not involve any endangered species.

## Author contributions

F.A.P, O.R. and N.N planned and designed the project. F.A.P and N.N. did the fieldwork and carried out the experiment. N.N and F.A.P. analysed the gene expression results. F.A.P. analysed the microbial data and interpreted the results. F.A.P and N.N wrote the article. O.R. helped during the fieldwork and supervised the project. O.R. supported the writing of the manuscript.

