## Supplementary material for "Parental fasting effects on offspring immune gene expression and gut microbiota in a species with male pregnancy (*Syngnathus typhle)*": Figure S1-S5

**2. Methods**

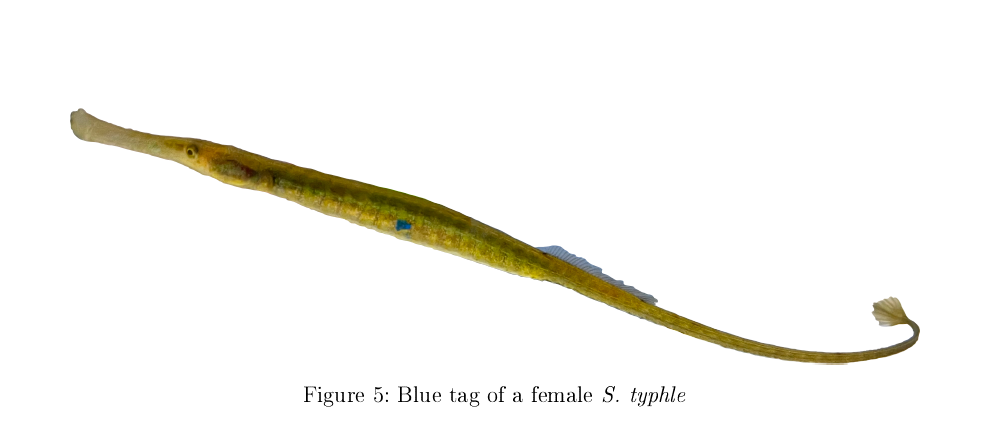

**Figure 1:** Example of a female *S. typhle* and the elastomer tagging.

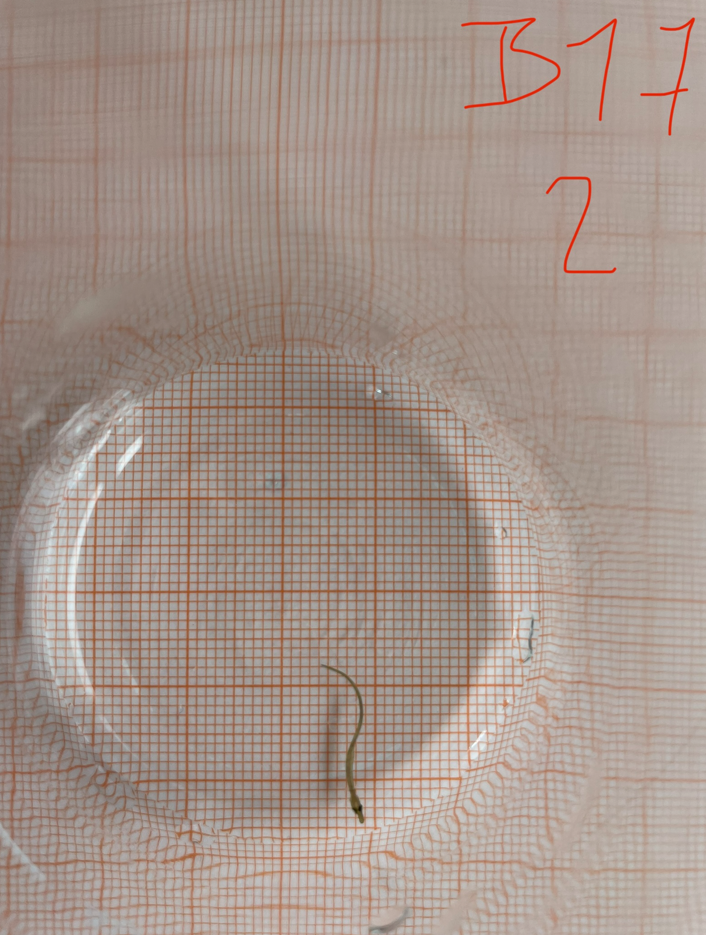

**Figure 2:** Example of size measurement of offspring *S. typhle* after birth using millimetre paper and then ImageJ.

**3. Results**

**3.1 Morphology**

For AL males, body length increased significantly from the start with 11.7 cm ± 0.44 to the end of the two months diet with 13.6 cm ± 0.43 (P < 0.0001), as did AL females (P < 0.0001) starting at 12.97 cm ± 0.89 and growing to 13.91 cm ± 0.79 after one month. In contrast, IF-treated individuals exhibited more modest growth: IF males from 13.0 cm ± 0.76 reaching 14.1 cm ± 0.78, and IF females from 12.93 cm ± 0.93 to 14.08 cm ± 0.87 (both P < 0.02) (Table S1 and S2_Morphology).

The initial size of AL males was 0.68 g ± 0.07, increasing to 1.03 g ± 0.09 after two months. In comparison, IF males started at 0.94 g ± 0.14 and reached 1.06 g ± 0.17. Female AL individuals began at 1.02 g ± 0.24 and grew to 1.13 g ± 0.19 after one month, while IF females started at 0.99 g ± 0.27 and reached 0.92 cm ± 0.23.

a)	b)

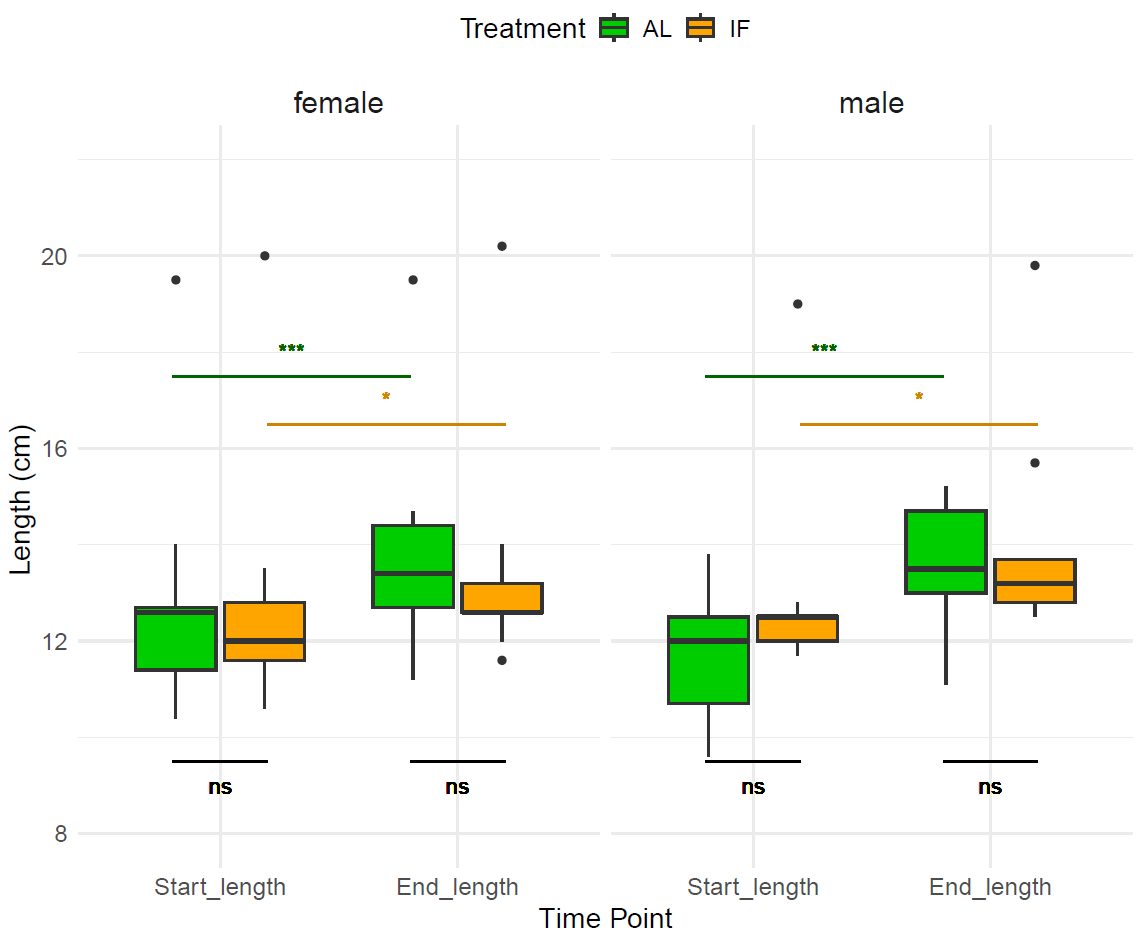

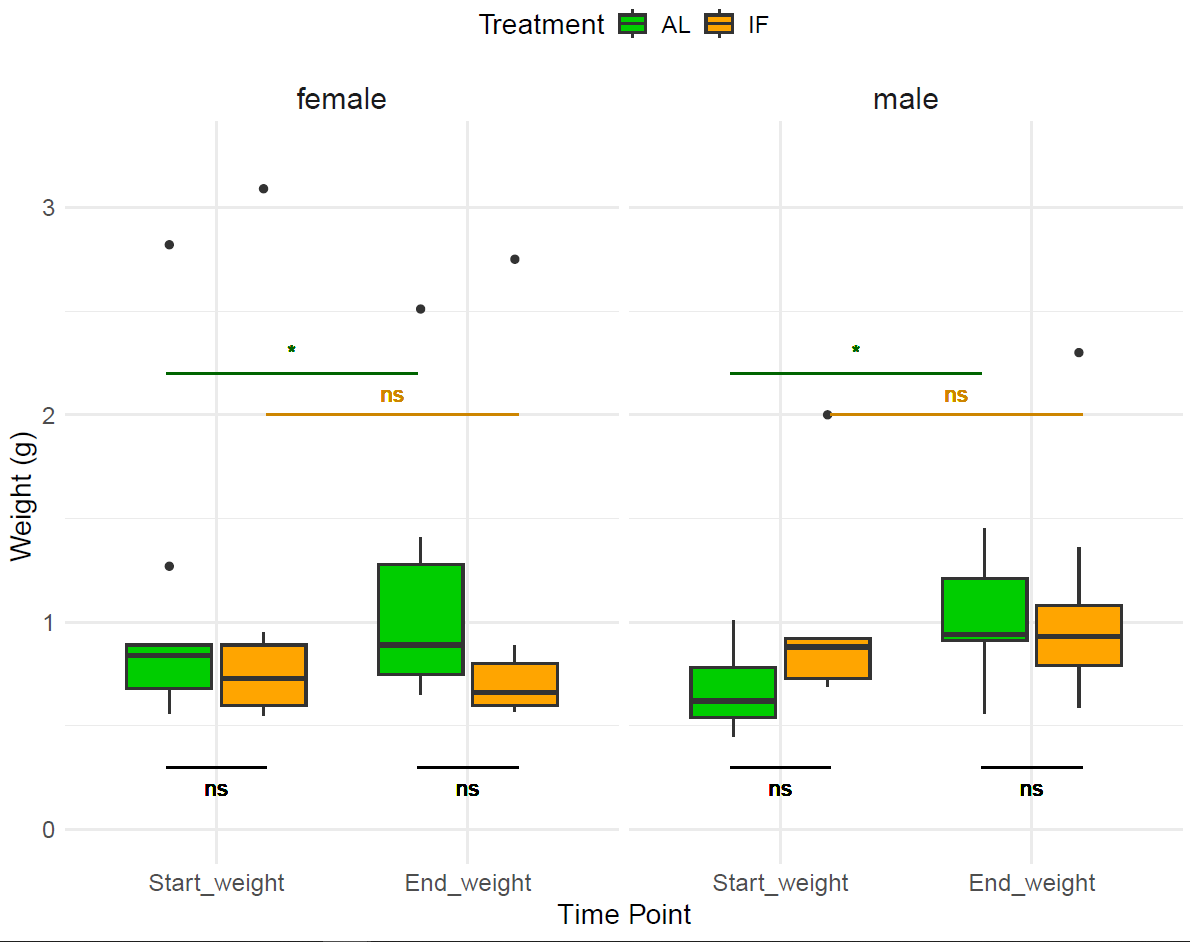

**Figure 3: Body length and weight of *S. typhle* parents**

a) The plot displays the body length of male and female pipefish before (Start_length) and after the treatment (End_length), with females shown on the left and males on the right. AL individuals are represented in green, and IF individuals are in orange. Two-way ANOVA results comparing treatment groups for both time points and sexes were not significant (ns). The linear mixed model results show significant changes in body size over time, with the IF group marked in orange (P < 0.02, *) and the AL group in green (P < 0.0001, ***). b) In this barplot we see the body weight of male and female pipefish before (Start_length) and after the treatment (End_length). Two-way ANOVA results comparing treatment groups for both time points and sexes were not significant (ns). The linear mixed model results show significant changes in body weight over time, for AL (P = 0.014, *), but not for IF (P = 0.86).

**3.2 Gene expression**

a)	b)

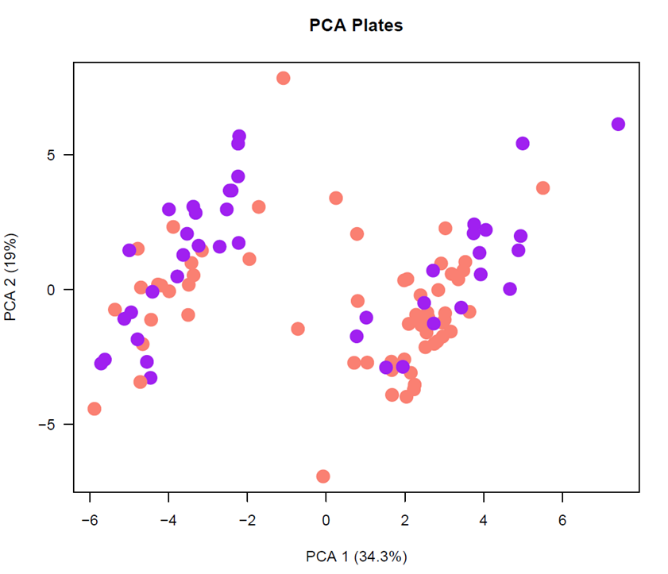

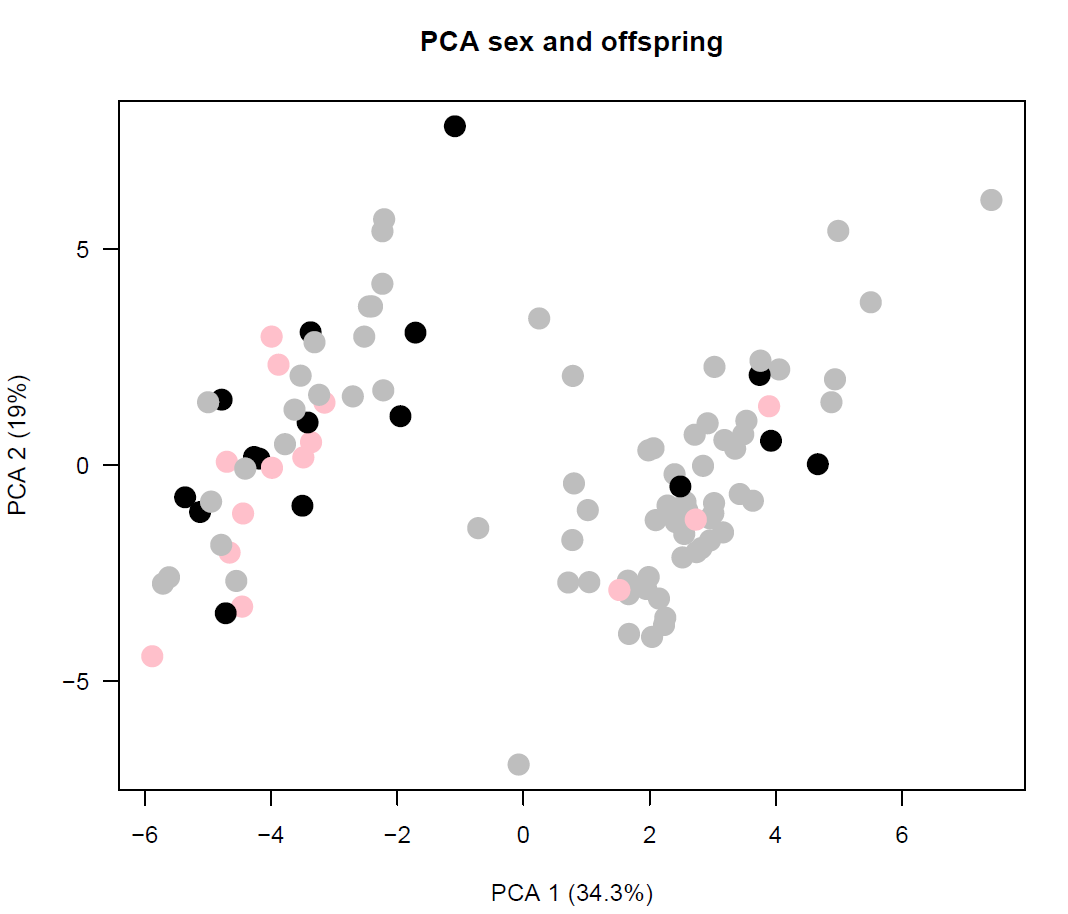

c)

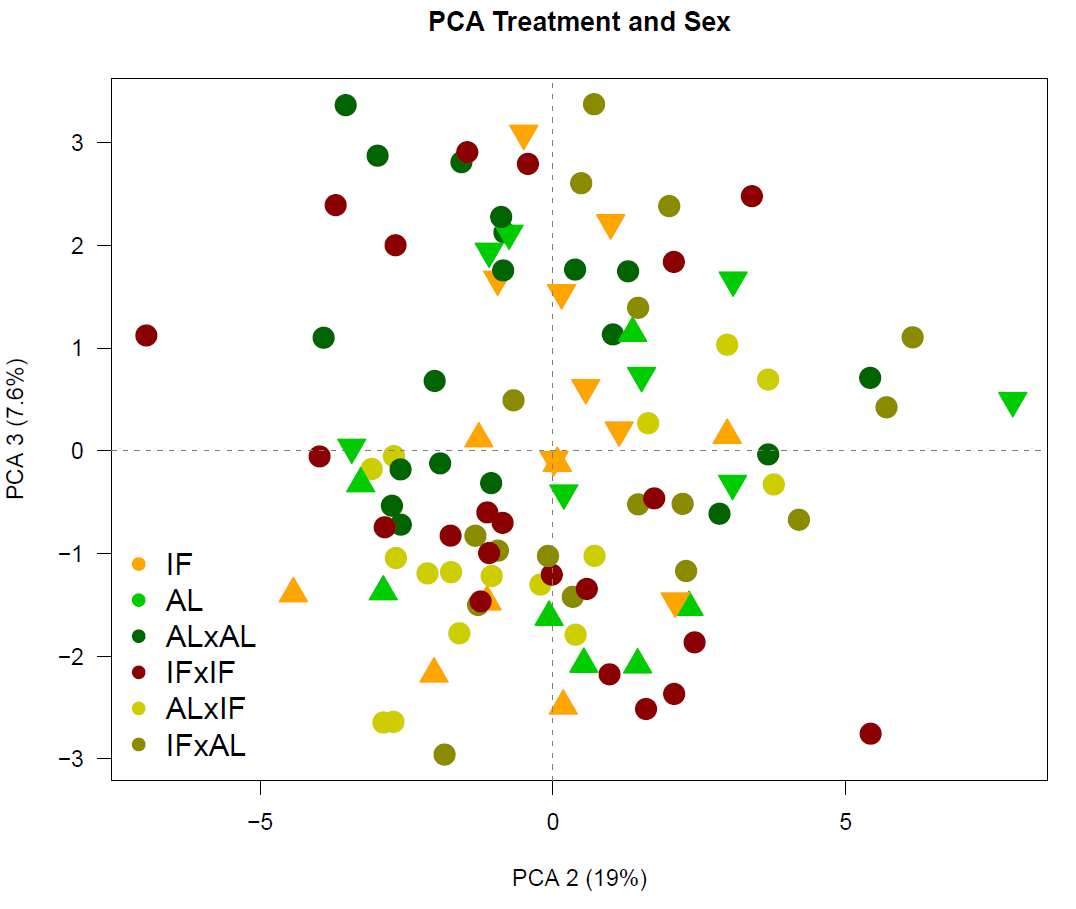

**Figure 4. Principal Component Analysis (PCA) of parental and offspring gene expression data obtained from Fluidigm analysis.**

Plots (a) and (b) represent PCA results for PC1 and PC2, with PC1 explaining 34.3% of the variance and PC2 accounting for 19%. In (a), dots are color-coded as purple or pink to differentiate between the two plate runs. A PERMANOVA on PC1 revealed a significant plate effect (P = 0.013), as well as a significant distinction between parents and offspring (P = 0.03). In (b), offspring data points are shown in gray, fathers in black, and mothers in light pink, indicating a trend for parental data to cluster towards the left of the PCA plot.

Plot (c) illustrates the PCA for PC2 and PC3, with PC3 explaining 7.6% of the variance. Here, data points are color-coded by treatment group, with triangles pointing down for females and up for males, distinguishing parental and offspring treatment groups. A PERMANOVA on PC3 showed a significant parent-offspring effect (P = 0.02) and a near-significant effect for treatment (P = 0.058).

a)	b)

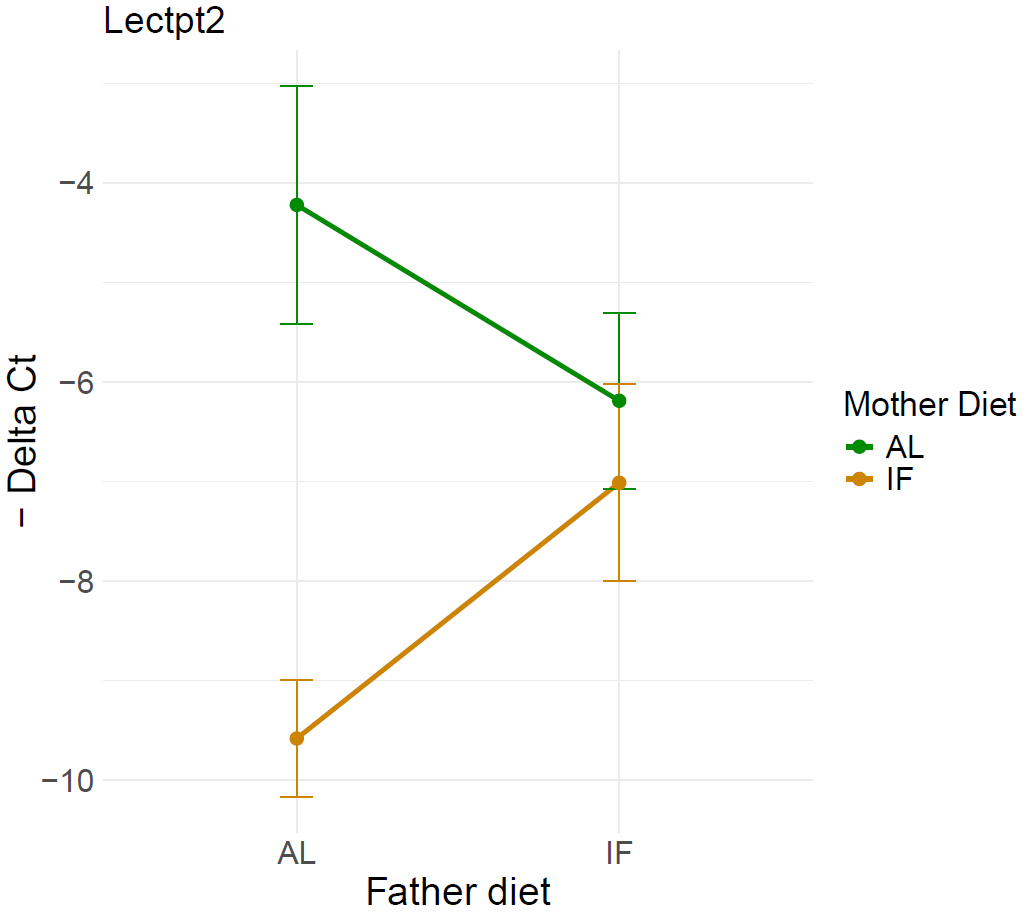

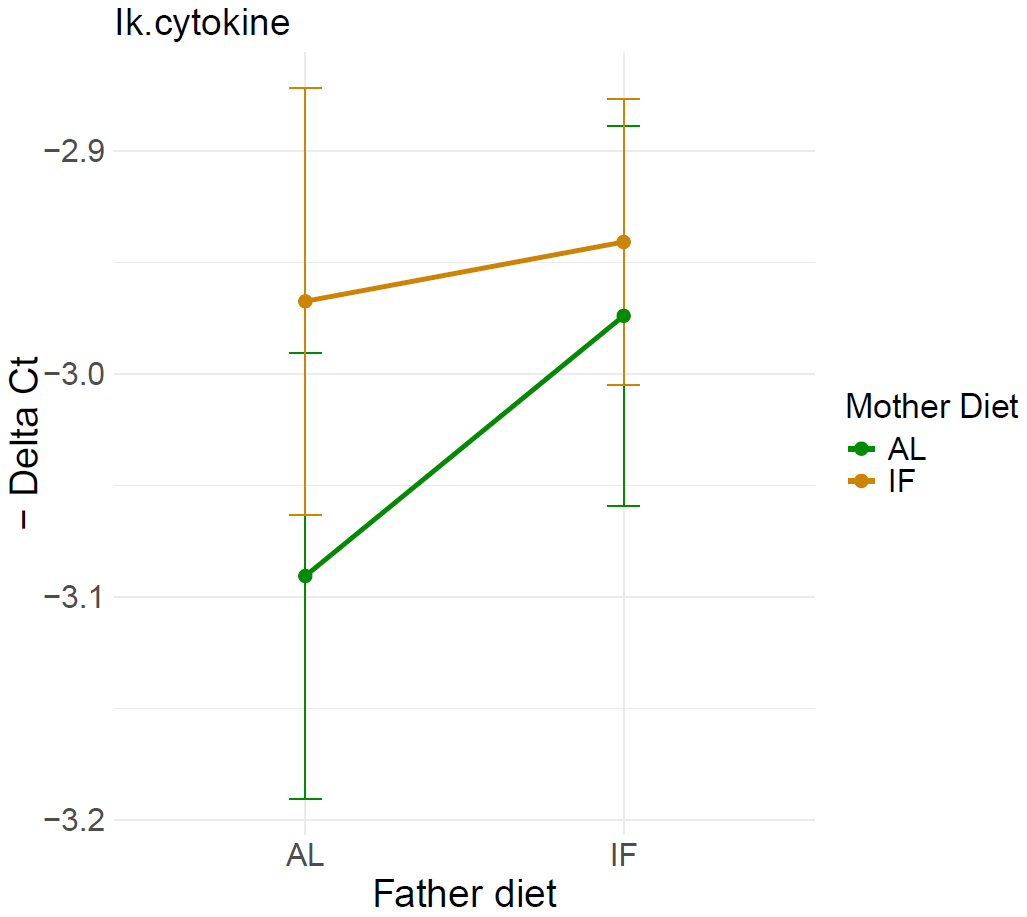

c)	d)

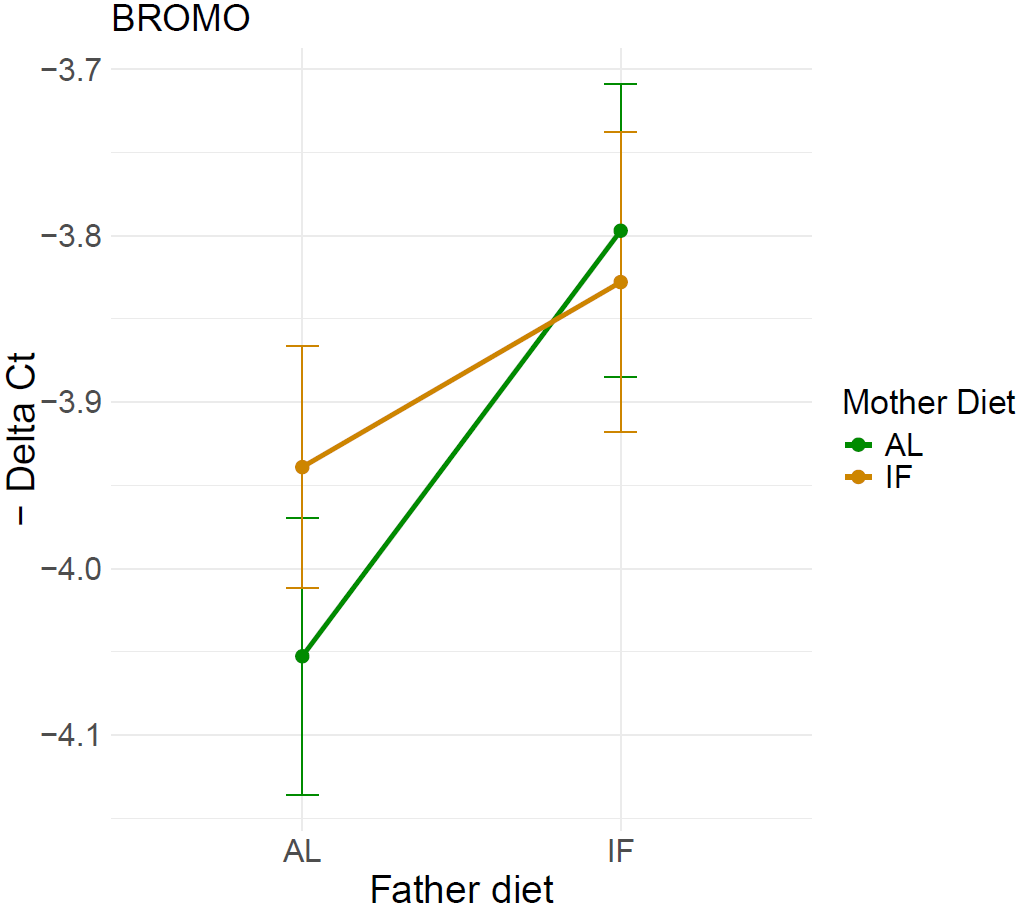

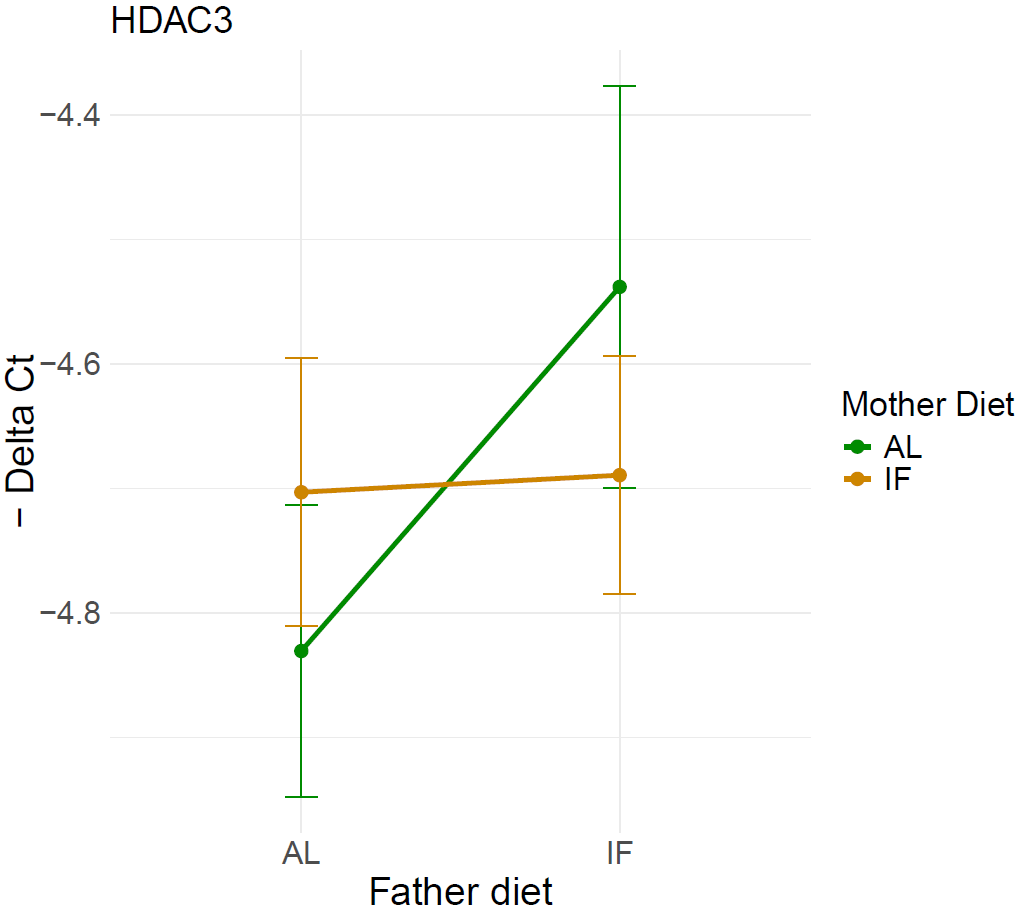

e)	f)

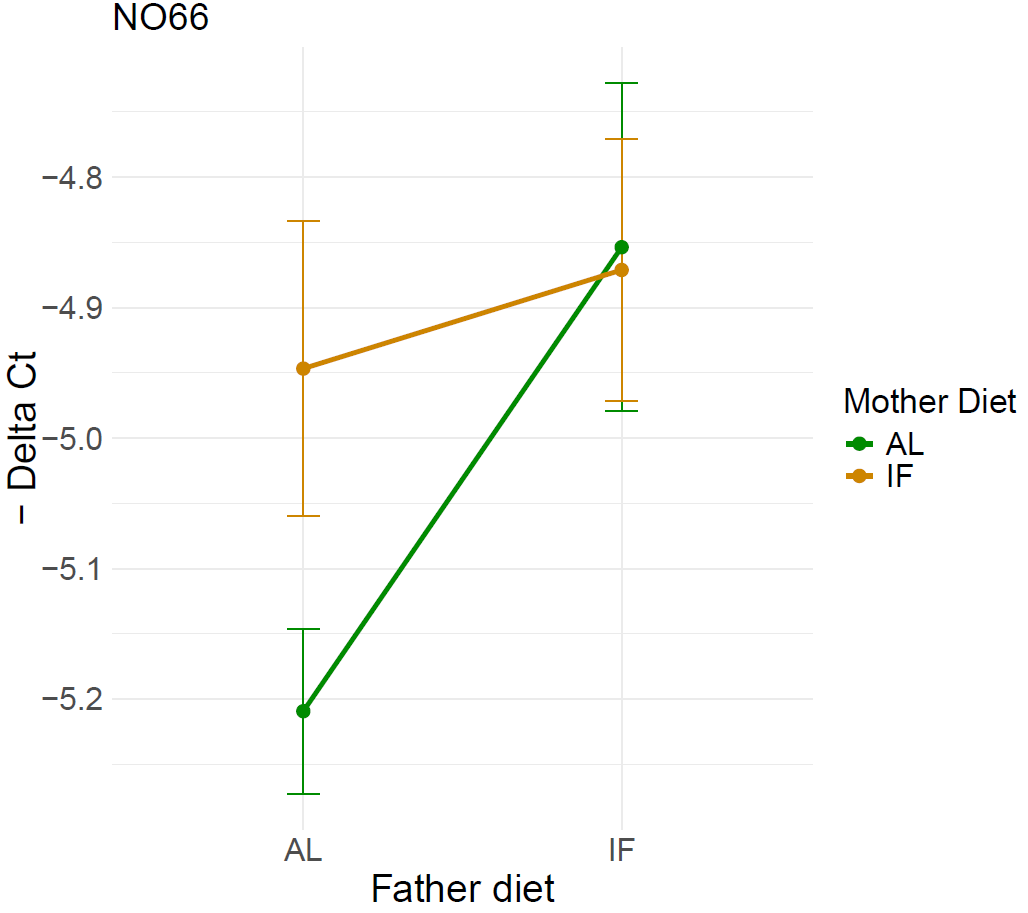

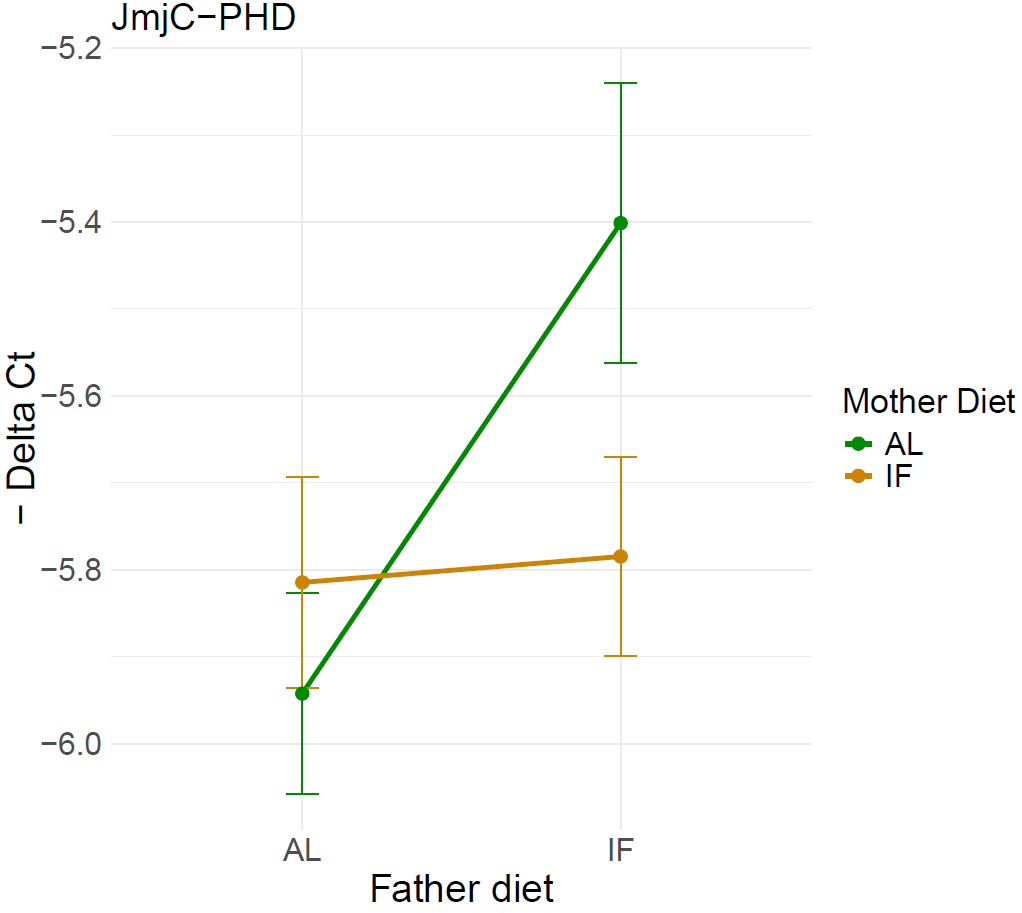

**Figure 5. Interaction plots of the most significantly differentially expressed genes in Syngnathus typhle offspring based on parental dietary treatments (additional details provided in the main manuscript).**

In these plots (a to f), the x-axis represents the father’s diet, while the colour coding (green for AL and orange IF) indicates the mother’s diet. The y-axis displays the negative Delta Ct values, which reflect the directionality of gene expression. Each plot illustrates the interaction between parental dietary treatments and their effects on offspring gene expression.
